# Metabolic adaptations of micrometastases alter EV production to generate invasive microenvironments

**DOI:** 10.1101/2024.05.12.593552

**Authors:** Michalis Gounis, America V. Campos, Engy Shokry, Louise Mitchell, Emmanuel Dornier, Nicholas Rooney, Sandeep Dhayade, Luis Pardo, Madeleine Moore, David Novo, Jenna Mowat, Craig Jamieson, Emily Kay, Sara Zanivan, Colin Nixon, Iain Macpherson, Saverio Tardito, David Sumpton, Karen Blyth, Jim C. Norman, Cassie J. Clarke

## Abstract

Altered cellular metabolism has been associated with acquisition of invasive phenotypes during metastasis. To study this, we combined a genetically engineered mouse model of mammary carcinoma with syngeneic transplantation and primary tumour resection to generate isogenic cells from primary tumours and their corresponding lung micrometastases. Metabolic analyses indicated that micrometastatic cells increase proline production at the expense of glutathione synthesis leading to a reduction in total glutathione levels. Micrometastatic cells also have altered sphingomyelin metabolism leading to increased intracellular levels of specific ceramides. The combination of these two metabolic adaptations alters small extracellular vesicle (sEV) production to drive generation of an invasive microenvironment. Indeed, micrometastatic cells shut-down Rab27-dependent production of sEVs and, instead, switch-on neutral sphingomyelinase-2 (nSM2)-dependent sEV release. sEVs released in a nSM2-dependent manner from micrometastatic cells, in turn, influence the ability of fibroblasts to deposit extracellular matrix which promotes cancer cell invasiveness. These data provide evidence that metabolic rewiring drives invasive processes in metastasis by influencing sEV release.

**Summary:** Breast cancer cells isolated from lung micrometastases have altered metabolism which influences extracellular vesicle production to generate invasive microenvironments.

## Introduction

Breast cancer can metastasise to various organs including the bone, liver, brain, and lung. Lung metastases are observed in approximately a quarter of patients with oestrogen receptor-positive metastatic breast cancer and in almost half of patients with the HER2-positive, oestrogen receptor-negative subtype ^1^. Thus, lung metastases represent a major contributor to breast cancer morbidity and mortality highlighting the need to understand the processes through which this cancer type colonises this organ. The first step in the metastatic cascade involves breaching the basement membrane and local invasion of the surrounding stroma, followed by entrance into the circulatory and/or lymphatic system and survival of tumour cells in these environments ^2, 3^. To metastasise to the lung, circulating cells must then extravasate into the lung parenchyma and seed small colonies (micrometastases) which must then grow to yield clinically detectable metastases. This last stage of the metastatic cascade poses a major bottleneck for development of clinically detectable metastases. Indeed, following extravasation, the majority of disseminated cancer cells are unable to grow in distant organ environments; some studies estimate that fewer than 0.02% of disseminated tumour cells can proceed to metastatic outgrowth ^4^. This inefficiency is likely due to vulnerability of extravasated tumour cells to elimination by the immune system, but also by challenges posed by the very different microenvironment that the tumour cells encounter there ^5^.

Successful metastasis requires that cancer cells rewire their metabolism to adapt to the various nutrient and metabolite profiles, and other factors such as oxygen tension and tissue stiffness, encountered at points on the metastatic cascade ^6^. There has been considerable focus on how such metabolic adaptations may support both growth and survival of metastasising cells. For instance, to survive oxidative insults encountered following detachment from the extracellular matrix (ECM) environment of the primary tumour site, disseminating cancer cells can increase pentose phosphate pathway activity to provide reducing equivalents for glutathione synthesis ^7, 8^. Moreover, evidence is accumulating that metabolic adaptations may support other cellular processes – such as cell migration/invasion and extracellular matrix production/remodelling - that cancer cells need to execute at various stages of the metastatic cascade. For example, stiff microenvironments increase activity of the creatine-phosphagen ATP-recycling system to power cytoskeletal dynamics during the invasive migration, and chemotaxis necessary to establish liver metastases of pancreatic ductal adenocarcinoma ^9^. Similarly related to invasive behaviour, transformation of mammary epithelial cells leads to metabolic stress which upregulates expression of the glutamate-cystine exchanger, xCT (SLC7A11) without affecting rates of glutaminolysis ^10^. Upregulated xCT leads to increased extracellular glutamate – which is manifest as increased levels of circulating glutamate in tumour-bearing individuals. Increased extracellular glutamate then activates metabotropic glutamate receptor signalling to promote intracellular trafficking of the pro-invasive matrix metalloprotease, MT1-MMP.

Having negotiated successful exit from the primary tumour, survival in the circulation, and extravasation, cancer cells must still adapt to the metabolic microenvironment of the metastatic target organ before they can form clinically detectable metastases. As discussed above, mammary cancer commonly metastasises to the lung and its metabolic microenvironment is very different from that of the mammary gland and the circulation. For instance, pyruvate is present in higher concentrations in lung interstitial fluid than in plasma ^11^ and, once metastasising cells have adapted to this, this nutrient supports several cellular processes necessary for metastatic outgrowth in this tissue. Indeed, metastasising mammary cancer cells adapt to using pyruvate: a) to provide α-ketoglutarate (αKG) to enable hydroxylation of proline residues in collagens ^12^; b) to increase activation of anabolic signalling through the mTOR pathway ^13^ and; c) to increase pyruvate carboxylase-dependent anaplerosis ^11^. Thus, adaptation to increased local pyruvate levels contributes to the ability of micrometastatic cells to condition the ECM niche – likely to increase their chances of survival and immune escape - and to provide building blocks and anabolic signalling for subsequent metastatic outgrowth.

Cancer aggressiveness is thought to be influenced by the primary tumour’s ability to alter the microenvironment in other organs to generate niches which render them receptive to metastatic seeding ^14^. Metastatic niche priming may involve mobilisation of elements of the innate immune system – such as macrophages and neutrophils – to suppress acquired immunity in metastatic sites ^15^. Also, alterations to the ECM which would be expected to support cancer cell survival, growth and invasiveness have been observed in metastatic target organs very early in metastasis and prior to the arrival and extravasation of cancer cells in those organs ^16^. Release of extracellular vesicles (EVs), particularly small EVs (sEVs) (also known as exosomes) from primary tumours is now established to assist with niche priming – often by altering ECM deposition in metastatic target organs. sEVs are generated within the endosomal system, and key components of the endosomal EV production and release pathway in cancer cells – particularly the Rab27 GTPases – are, therefore, key to priming of metastatic niches in both the liver and lung in mammary cancer ^17^.

Evidence is accumulating that altered metabolic landscapes can influence sEV production, and this is now thought to be a mechanism through which cells may communicate information concerning the state of their metabolism to other cells. For instance, in obesity, metabolically-stressed/damaged hepatocytes secrete sEVs which can communicate with liver stellate cells to promote ECM deposition thus increasing the fibrosis associated with fatty liver ^18^. Also, mitochondrial stress in mammary cancer cells activates the PINK1 kinase which leads to packaging of mitochondrial DNA (mtDNA) into sEVs ^19^. These mtDNA-containing sEVs can then drive invasive behaviour in other cells via activation of a toll-like receptor signalling. The metabolic rewiring occurring in disseminated cancer cells as they adapt to the microenvironment of other organs may, therefore, modulate release of factors such as sEVs that are able to generate ECM niches that permit metastatic outgrowth.

Here we have deployed a genetically-engineered mouse model (GEMM) of mammary cancer to generate cells from very early lung micrometastases which are isogenically-paired with their corresponding primary tumour cells. This approach has enabled in-depth characterisation of the metabolic adaptations made by cancer cells very early in metastatic seeding, and a description of how these influence release of sEVs that can influence ECM deposition to generate a microenvironment conducive to subsequent invasive growth.

## Results

### Generation and characterisation of micrometastatic cells

Orthotopic transplantation models can be used to generate organ-specific isogenic metastatic cell lines to study adaptations made by breast cancer cells as they move from their primary tumour site to metastatic target organs, such as the lung ^20^. As we are interested in how metabolic adaptations might sculpt the early metastatic microenvironment of mammary cancer, we sought to establish cultures of cells that have recently relocated from their primary site in the mammary gland to the lung. To do this, we re-introduced cells derived from primary tumours in the MMTV-*PyMT* model of mammary cancer (maintained in the FVB mouse strain) ^21^ – termed parental (P) cells - into the 4^th^ mammary fat pad of FVB mice (Fig. 1A). We allowed these to grow into tumours of 8-10mm in diameter, resected them from the mammary fat pad, placed them into culture, and thereafter refer to these as fat pad (FP) cells. Post-resection of the primary tumour, mice were maintained for approximately 1 month to allow metastases to seed. Mice were then sacrificed, their lungs minced and placed into culture. We then cultured the *PyMT*-expressing cancer cells which grew-out from these lung homogenates – thereafter referred to as micrometastatic (M) cells. We generated two series of these cells (P, FP, M and P’, FP’ and M’ respectively) using independent MMTV-*PyMT* mice. All cells in these series expressed the *PyMT* antigen and E-cadherin (not shown), confirming that they were tumour cell-derived and of epithelial origin. Moreover, cells from primary tumours (P/P’ and FP/FP’) and micrometastases (M/M’) all grew at similar rates (Fig. 1B; S1A). However, micrometastatic cells (M/M’) were significantly more migratory than their primary tumour counterparts (P/P’ or FP/FP’) as determined by transmigration towards a gradient of fibronectin and serum (Fig. 1C; S1B) or by measuring penetration of cancer cells into ‘organotypic’ plugs of collagen which had been preconditioned with telomerase-immortalised dermal fibroblasts (TIFs) ^22^ (Fig. 1D, E; S1C).

**Fig. 1.**
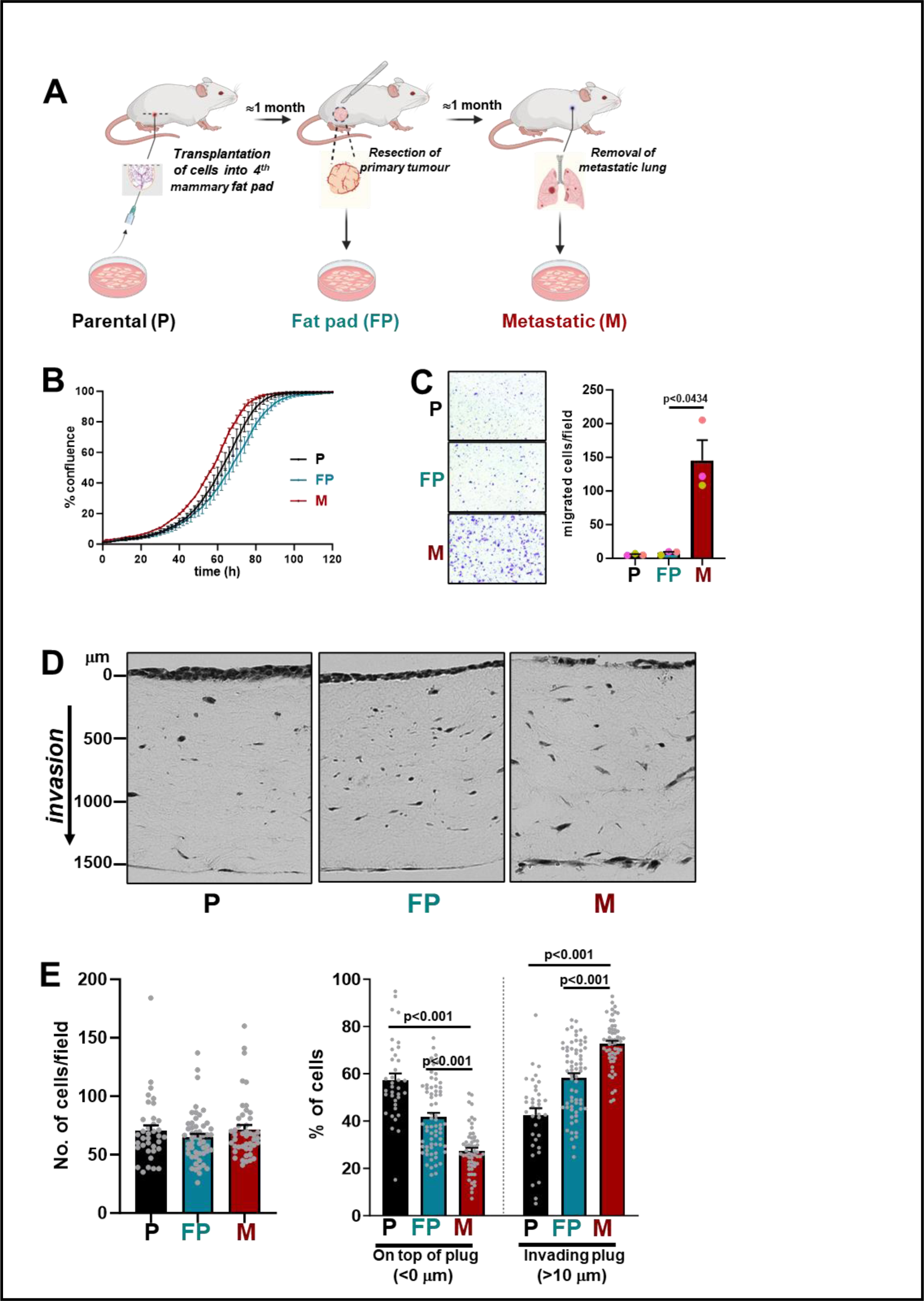
Cells from lung micrometastases display increased invasiveness. **(A)** Parental (P) cell lines were established from mammary tumours spontaneously arising in MMTV-*PyMT* female mice. These cells were then transplanted into the 4^th^ mammary fat pad of syngeneic mice (FVB/N), and tumours grown to a defined size (8-10mm). Tumours resected from the mammary fat pad were used to establish ‘fat pad’ (FP) cell lines. Following tumour resection, mice were maintained for sufficient time (1 month) to allow seeding of micrometastases in the lung. Lungs were then removed, and *PyMT*-positive ‘micrometastatic’ (M) cell lines were established from lung homogenates. **(B)** P, FP, or M cells (as described in (A)) were plated onto 6-well dishes and their growth determined using the IncuCyte ZOOM live-cell imaging system. Values are mean ± SD, n=3 technical replicates/cell line). **(C)** P, FP, or M cells were seeded into the upper chambers of Transwells (8µm pore size) and transmigration over a 2 hr period toward a gradient of serum and fibronectin (applied to the lower) chamber was determined. Representative images (left panels) and quantification (right panels) of the number of cells adherent to the upper surface of the lower chamber are displayed. Values are mean ± SEM, n=3 independent experiments; data were analysed using a paired t-test. **(D & E)** P, FP or M cells were plated onto plugs of rat tail collagen that had previously been conditioned by telomerase-immortalised dermal fibroblasts (TIFs) for 4 days. Cancer cells were allowed to invade pre-conditioned plugs for 7 days followed by fixation and visualisation of cells using H&E. Representative images (D) and quantification of the total number (E, left graph) and % (E, right graph) of cells remaining on top of or invading collagen organotypic plugs to a depth of >10µm are displayed. Values are mean ± SEM, n=36-52 fields of view/cell line (2 biological replicates for FP and M cells and 1 biological replicate for P cells), data were analysed by mixed effects ANOVA with Tukey’s multiple comparison test.

### Micrometastatic (but not macrometastatic) cells increase proline synthesis at the expense of glutathione production

As we were interested in how metastasising mammary cancer cells might rewire their metabolism during the early stages of seeding the lung, we compared how the cells we have established from MMTV-*PyMT* primary tumours (P & FP) and their micrometastases (M) altered the profile of extracellular metabolites over a 24hr period. Although most metabolites were either consumed (e.g., glucose and glutamine) or secreted (e.g., lactate and alanine) at similar rates in all cells, one metabolite (proline) had markedly altered consumption/secretion dynamics between primary tumour-derived (P and FP) and micrometastatic (M) cells (Fig. 2A). Strikingly, proline was consumed by primary tumour-derived cells, whereas micrometastatic cells secreted this amino acid (Fig. 2A). To determine whether increased proline secretion by metastatic cells might manifest in vivo, we profiled the circulating metabolome of MMTV-*PyMT* mice and healthy age-matched control animals. We determined the lung metastatic burden of tumour-bearing mice and subdivided them into those that did (Mets) and did not (No mets) display lung metastases, as determined by retrospective histological examination (not shown). This indicated that MMTV-*PyMT* mice bearing lung metastases displayed significantly higher levels of circulating proline than MMTV-*PyMT* mice that had extensive primary tumour growth in the mammary gland, but no metastases (Fig. 2B). By contrast, other amino acids (such as asparagine and serine) whose levels did not differ between conditioned media from P, FP or M cells (Fig. 2A), were similar in mice that did and did not bear metastases (Fig. 2B). This result from a mouse model of metastatic mammary cancer encouraged us to compare levels of proline in the circulation of healthy volunteers and patients with metastatic breast cancer. Circulating proline (but not asparagine or serine) was significantly elevated in the plasma of patients with metastatic breast cancer indicating that this amino acid may represent a potentially useful biomarker for metastases in breast cancer (Fig. 2C).

**Fig. 2.**
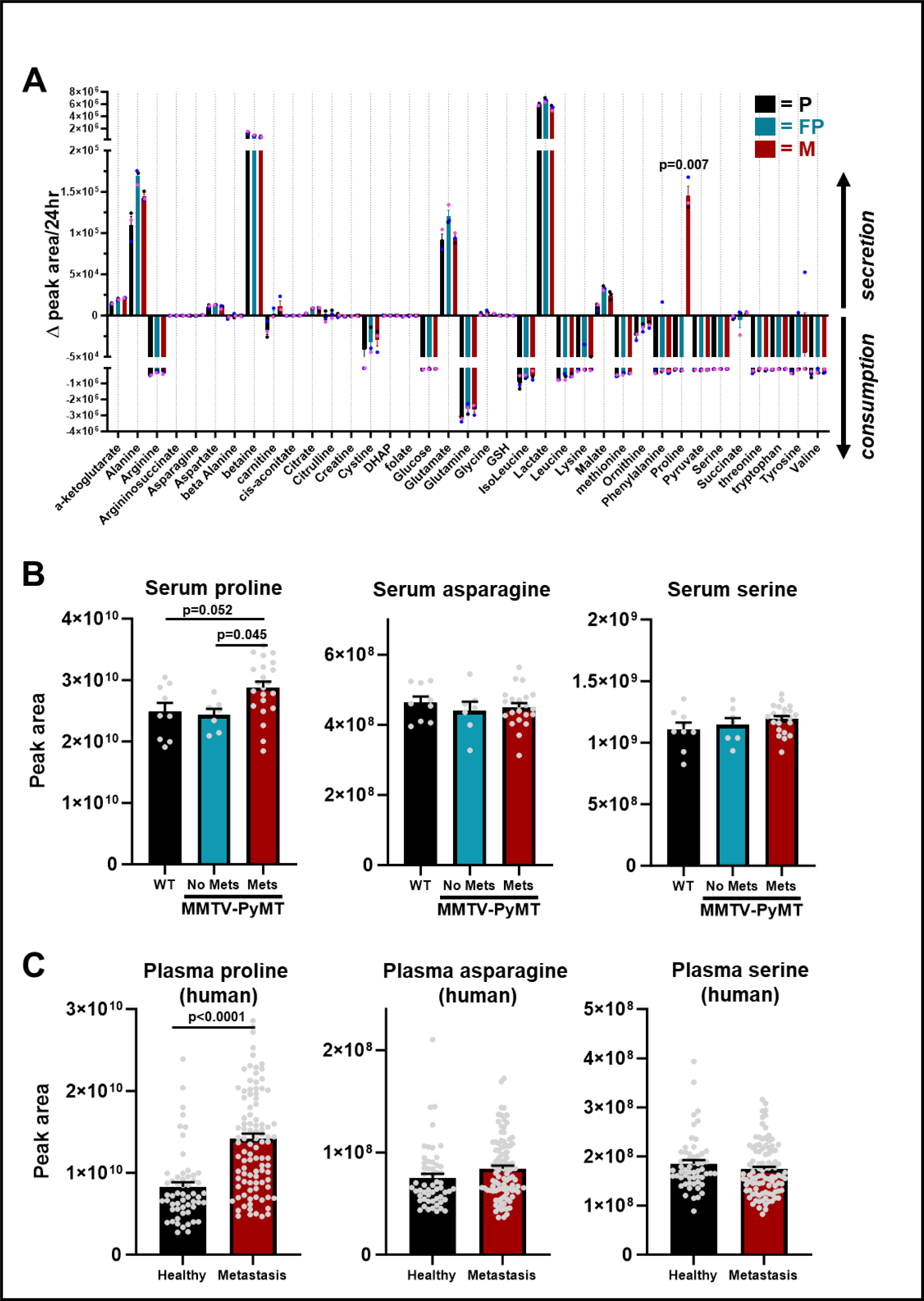
Micrometastatic cells secrete proline. **(A)** Parental (P), fat pad (FP) or micrometastatic (M) cells were plated onto six-well dishes and incubated at 37°C for 24 hr. Conditioned medium was aspirated, and levels of the indicated metabolites determined using LC-MS-based metabolomics. Data are expressed as the difference between metabolite levels detected in cell-conditioned medium and those in medium incubated at 37°C in the absence of cells (Δ peak area). Thus, positive values indicate production and release of a metabolite by cells, whereas negative values indicate consumption of that metabolite during the 24 hr period. Values are mean ± SEM, n=3 independent experiments, data were analysed by two-way ANOVA with Tukey’s multiple comparison. **(B)** MMTV-*PyMT* and non-tumour bearing FVB control mice were culled at 11-14 weeks of age, blood was collected by puncture of the posterior vena cava, sera prepared by centrifugation and levels of the indicated metabolites determined using LC-MS. Lungs of MMTV-*PyMT* mice and non-tumour bearing FVB control mice (n=9 mice) were assessed by histology and categorized according to the presence (Mets; n=22 mice) or absence (no Mets; n=7 mice) of lung metastases. Values are mean ± SEM, data were analysed by ordinary one-way ANOVA with Sidak’s multiple comparison. **(C)** LC-MS metabolomics was used to determine levels of the indicated metabolites in plasma collected from metastatic breast cancer patients (n=99) and healthy volunteers (n=56). Values are mean ±SEM, data were analysed using unpaired Student’s t-test.

We then proceeded to look for intracellular polar metabolites that were elevated within micrometastatic cells by comparison with their primary tumour counterparts. Only one metabolite, proline, was increased in both series of micrometastatic cells (M & M’) by comparison with their primary tumour counterparts (P & P’; FP & FP’) (Fig. 3A, B S2A, B), consistent with our finding that micrometastatic cells release more proline, and that this amino acid is specifically elevated in the circulation of mice with lung metastases. We detected only one metabolite - glutathione (in both its reduced (GSH) and oxidized (GSSG) forms) - that was significantly decreased in micrometastatic cells (M & M’) by comparison with cells from primary tumours (P & P’; FP & FP’) (Fig. 3A, B; S2A, B). To further confirm these alterations to GSH levels, we deployed the thiol alkylating reagent N-ethylmaleimide (NEM) - a cell-permeable agent that reacts with GSH (but not GSSG), prior to metabolite extraction ^23, 24^. This indicated that GS-NEM (the glutathione adduct of NEM) levels were decreased in NEM-treated micrometastatic cells, indicating that these cells display genuinely decreased levels of GSH (Fig. 3C). *De novo* glutathione biosynthesis involves two ATP-dependent enzymatic steps, where ligation of cysteine to glutamate is catalysed by glutamate-cysteine ligase (GCL) the rate-limiting enzyme for glutathione biosynthesis to form γ-glutamylcysteine ^25^. Subsequently, glycine is added to this dipeptide by glutathione synthetase (GSS) to form the tripeptide, GSH. Although cysteine levels remained unchanged, we found that levels of γ-glutamylcysteine were decreased in micrometastatic cells (Fig. 3C). To determine whether expression of glutathione biosynthetic enzymes might explain these reductions in GSH and GSSG levels, we used qPCR to quantify the expression of *Gclc*, encoding the catalytic subunit of GCL. *Gclc* expression was suppressed in micrometastatic cells (M), by comparison with cells from primary tumours (P and FP), indicating that suppression of GCL activity may be one reason underlying decreased glutathione levels in micrometastatic cells (Fig. 3D, left graph). By controlling intracellular levels of cystine and cysteine, the glutamate-cystine antiporter, xCT can also influence glutathione levels ^25^. xCT is upregulated in various human cancers and previously published work has highlighted the ability of *PyMT*-derived primary tumour cells to release glutamate via increased xCT expression ^10^. qPCR indicated that micrometastatic cells express almost 50% less xCT (*Sclc7a11*) than their primary tumour-derived counterparts (Fig. 3D, right graph). To further assess the (patho)-physiological relevance of our finding, we used RNA *in situ* hybridisation to compare xCT expression in primary tumours and the micrometasases present in the lungs of matched MMTV-*PyMT* mice. This indicated that primary tumours express substantial amounts of xCT, whereas this was significantly decreased in lung micrometastases from the same animals (Fig. 3E). Importantly, the surrounding lung parenchyma expressed more xCT than the micrometastases indicating that in situ hybridisation was effective in this tissue. These data indicate that xCT is downregulated in lung micrometastases by comparison with primary tumour cells and that this, in combination with the reduction in GCL levels may lead to decreased glutathione in micrometastatic cells.

**Fig. 3.**
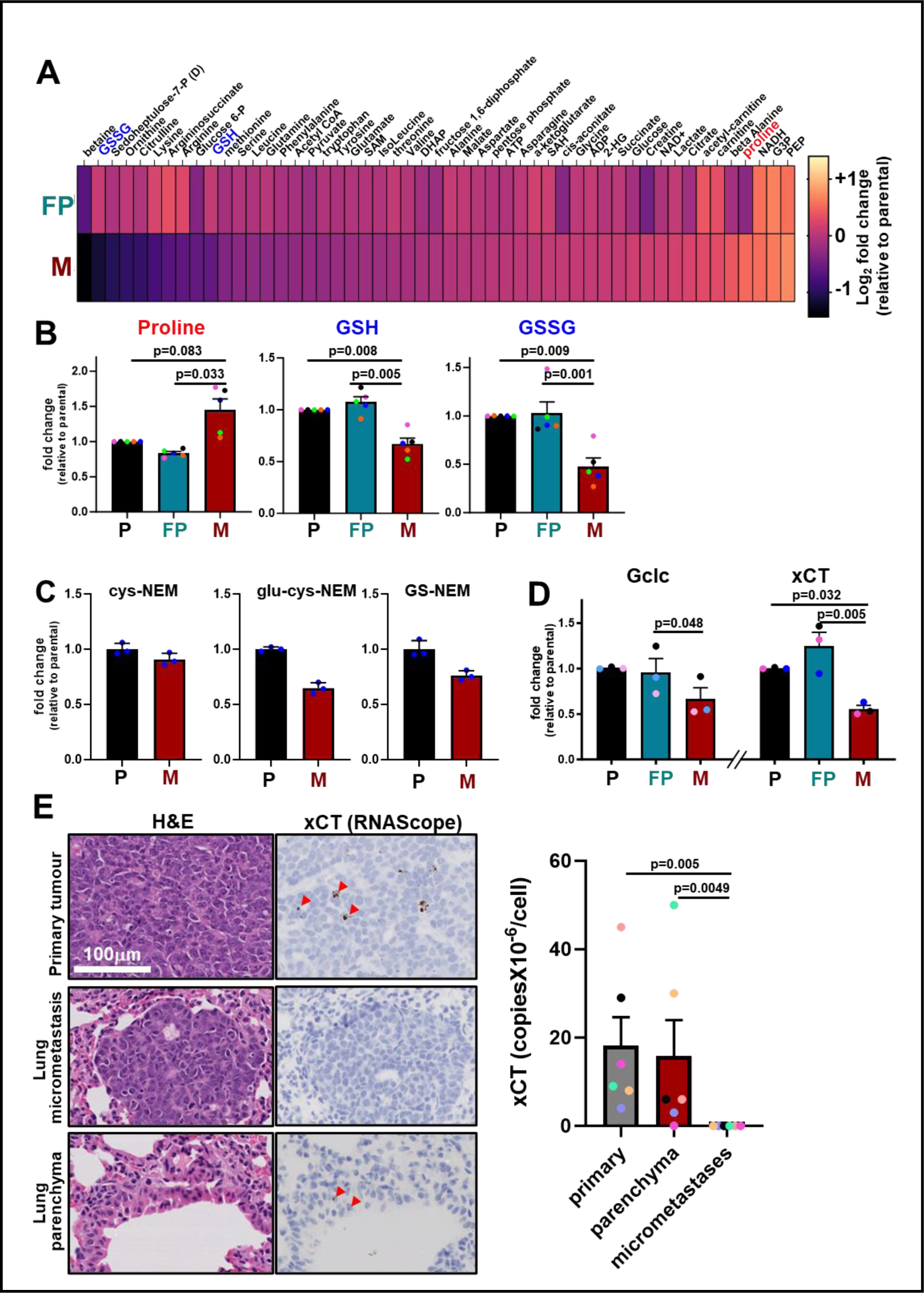
Proline and glutathione metabolism is altered in micrometastatic cells. **(A, B)** Parental (P), fat pad (FP) or micrometastatic (M) cells were cultured as for figure 2A and levels of intracellular metabolites determined using LC-MS-based metabolomics. The abundance of intracellular metabolites in fat pad (FP) and micrometastatic (M) cells were normalized to cell number and expressed relative to levels of the same metabolites in parental (P) cells. For the heatmap in (A), values are log_2_-fold changes, and for (B) values are mean fold-change, data were analysed by repeated measures one-way ANOVA with Tukey’s multiple comparison test, n=5 independent experiments. (GSH, reduced glutathione; GSSG, oxidised glutathione; SAM, S-adenosyl methionine; DHAP, dihydoxyacetone phosphate; G3P, glycerol-3-phosphate; PEP, phosphoenolpyruvate, S-adenosylhomocysteine). **(C)** Parental (P) and micrometastatic (M) cells were cultured as for figure 2A and lysates derivatised with N-ethylmaleimide (NEM). NEM adducts of cysteine (cys-NEM), γ-glutamylcysteine (glu-cys-NEM) and reduced glutathione (GS-NEM) were then detected using LC-MS and plotted as fold change relative to parental cells, n=1, values represent mean ±SD (3 technical replicates are shown). **(D)** Cells were cultured as for figure 2A and levels of the mRNAs encoding glutamate-cysteine ligase catalytic subunit (*Gclc*; **left graph**) and *Slc7a11* (xCT; **right graph**) were determined using qPCR. Values were normalised to *ARPP0* mRNA levels and expressed as fold change relative to parental (P) cells. Values are mean ± SEM, n=3 independent experiments, data were analysed using paired t-test. **(E)** Mammary tissue and lungs were resected from MMTV-*PyMT* mice at primary tumour endpoint (tumour size 15 mm). Tissues were formalin fixed, paraffin embedded, and sectioned for histological examination. Expression of the mRNA encoding xCT/*Slc7a11* was visualized using in situ hybridisation (RNAScope) and sections were counterstained with H&E. Sections representing primary mammary tumours, their matched lung micrometastases and surrounding lung parenchyma are displayed. Red arrowheads highlight the brown dots that indicate hybridisation with the xCT probe, bar 100μm. Quantification of the average optical density of the xCT probe in stained tissue sections was achieved using HALO software, values are mean ± SEM, Kruskal-Wallis with Dunn’s multiple comparisons test, n=6 mice/group.

In addition to culturing cells from micrometastases (M & M’) we have dissected larger/frank metastases from the lung of MMTV-*PyMT* tumour-bearing mice and established cultures from these (maM’ cells). Metabolomic analysis indicated that cells cultured from frank metastases display similar GSH, GSSG and proline levels to their primary tumour-derived counterparts (Fig. S2C, D). Thus, adoption of a metabolic profile in which proline and glutathione levels are respectively elevated and suppressed is a feature that is unique to micrometastatic cells and was not apparent in cells derived from a larger/frank metastasis.

Because glutamate is a precursor of both glutathione and proline, we hypothesised that levels of these metabolites might move in opposing directions when cancer cells form micrometastases, perhaps due to competition for carbons from this shared precursor. We tested this by performing metabolic tracing using ^13^C_5_-labelled glutamine. In cells from primary tumours (P) glutamine-derived carbons were found predominantly in glutathione and Krebs cycle intermediates, such as α-ketoglutarate (αKG) (Fig. 4A). Conversely, in micrometastatic cells (M), glutamine-derived carbons were present in increased levels in both the cellular and secreted pools of proline (Fig. 4A). To directly test whether the glutathione and proline synthesis pathways compete for these carbons we treated micrometastatic cells with an inhibitor of pyrroline-5-carboxylate reductase (PYCR) a key enzyme in the proline synthesis pathway ^26, 27^. As expected, the PYCR inhibitor (PYCRi) opposed synthesis (and secretion) of proline from glutamine (Fig. 4B). In addition to inhibiting synthesis of proline from glutamine, this inhibitor led to significant dose-dependent increases in the flux of glutamine-derived carbons towards αKG and glutathione synthesis in micrometastatic cells (Fig. 4B).

**Fig. 4.**
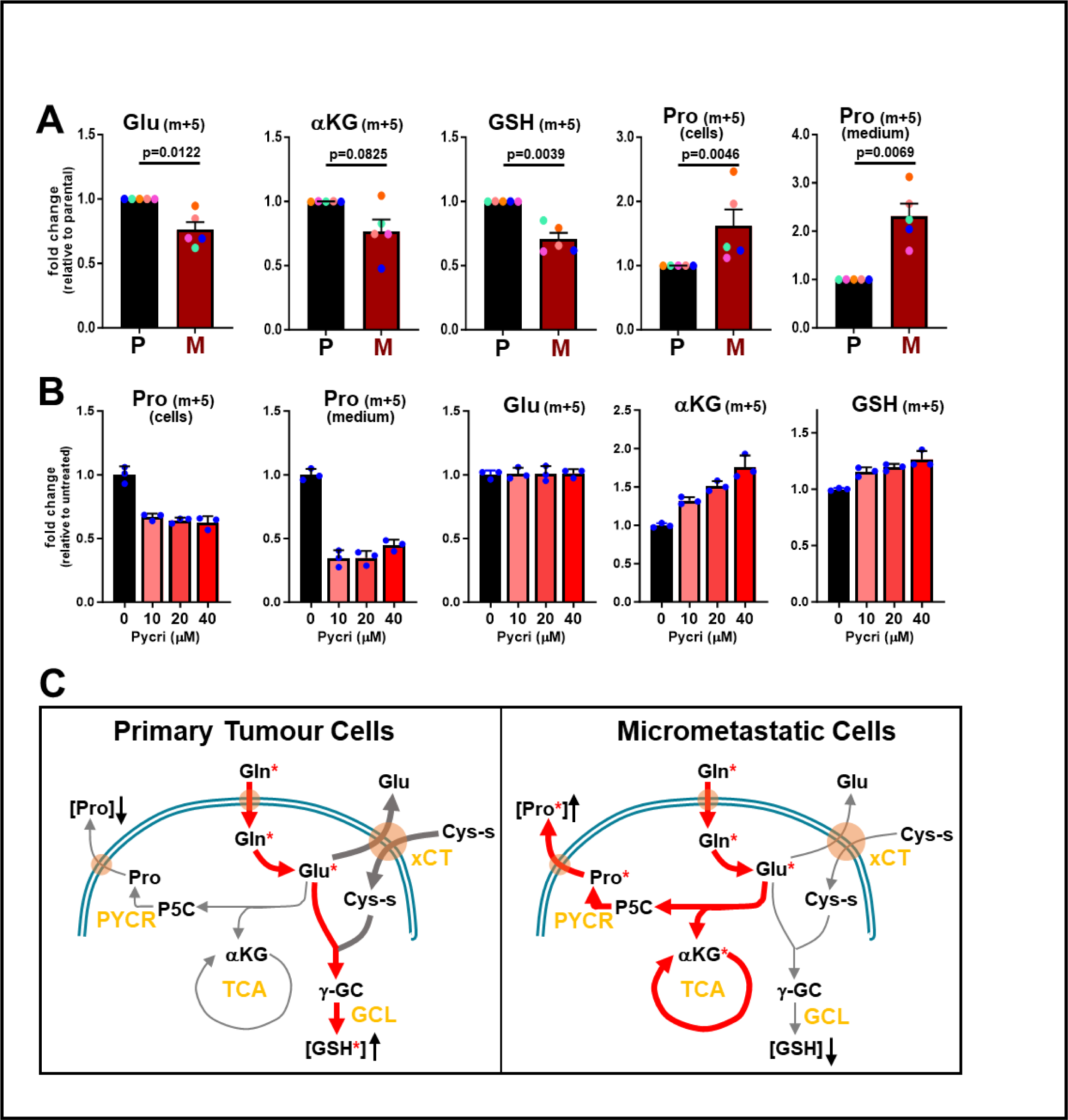
Proline and glutathione synthesis pathways compete for glutamine-derived carbons in micrometastatic cells. **(A, B)** Parental (P) or micrometastatic (M) cells were cultured for 24h in the presence of ^13^C_5_-labelled glutamine (A). Additionally, micrometastatic (M) cells were cultured with ^13^C_5_-labelled glutamine in the presence of the indicated concentrations of PYCR inhibitor (PYCRi) for 24 hr (B). The presence of ^13^C-labelled glutamine-derived carbons in the indicated metabolites was determined using LC-MS. Values are mean ± SEM, n=5 (for (A)), data were analysed by paired t-test. n=1 (3 technical repeats) (for (B)). **(C)** Schematic representation of the utilisation of glutamine-derived carbons (denoted with red asterisks) in *PyMT*-derived primary tumour (P) and micrometastatic (M) cells. Glutamine-derived carbons are primarily used for glutathione production in primary tumour cells, while micrometastatic cells re-route these carbons towards proline synthesis at the expense of glutathione production (Cys-s, cystine).

Taken together, these data indicate that mammary cancer cells that have left the primary tumour and are in the early stages of seeding metastases in the lung increase the glutamine-dependent production of proline, which is then exported from the cells, and that this occurs at the expense of glutathione synthesis (Fig. 4C). Moreover, this situation is likely to be transient because our data indicate that the balance of proline and glutathione synthesis returns to levels similar those observed in primary tumours as micrometasases evolve into frank metastatic outgrowth.

### Decreased glutathione synthesis is associated with increased sEV release

We and others have established that sEV release from primary tumours contributes to priming of the lung metastatic niche ^14, 16^. Moreover, we have shown that sEVs from tumours with high metastatic potential – such as those that have acquired particular gain-of-function mutations in p53 - can generate pro-invasive microenvironments which are associated with tumour dissemination and metastatic seeding ^16^. We were, therefore, interested in measuring sEV release from micrometastatic mammary cancer cells and how this might be related to their altered metabolism. We used differential centrifugation to purify EVs from cell-exposed culture medium and this indicated that micrometastatic cells (M & M’) released significantly more CD63-positive sEVs than their primary tumour counterparts (P & P’; FP & FP’) (Fig. 5A, B; S3A, B). We then used buthionine sulfoximine (BSO), an irreversible inhibitor of GCL, to interrogate a causal relationship between decreased glutathione levels and increased sEV release. BSO has been extensively used to evoke oxidative stress by depletion of GSH reserves ^28, 29^. By careful titration we identified concentrations of BSO (0.625 – 2.5 μM) that decreased total glutathione (GSSG and GSH) levels in cells from primary tumours to those observed in micrometastatic cells (approx. 40-50% reduction) (Fig. 5D). Moreover, we were able to decrease GSH levels in a manner that was reproducibly stable for 48hr (not shown) to allow for collection of sEVs, and that did not compromise cell growth/viability (not shown) or detectably disturb levels of other cellular metabolites (Fig. 5C). Restricting glutathione synthesis in this way led to significantly increased release of CD63-positive sEVs from cells derived from primary tumours (P) (Fig. 5E). This indicates that the increased capacity of micrometastatic cells to release sEVs is likely to be, at least in part, due to the decreased synthesis of glutathione observed in these cells.

**Fig. 5.**
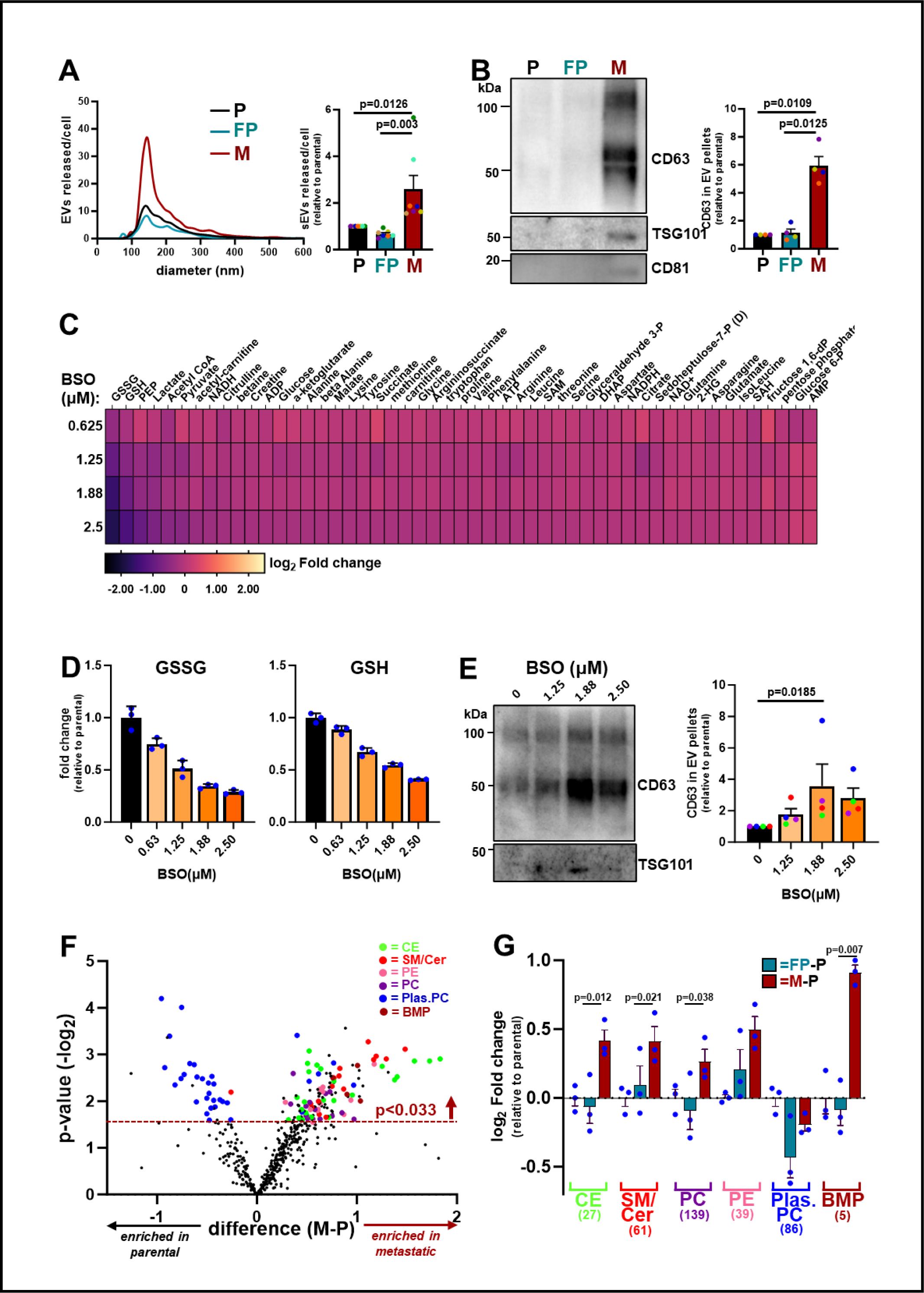
Decreased glutathione synthesis in micrometastatic cells is associated with increased sEV release. **(A, B)** Conditioned media were collected from parental (P), fat pad (FP) and micrometastatic (M) cells and sEVs were purified from these using differential centrifugation. The size distribution and number of EVs were analysed by nanoparticle tracking (A) and the CD63, TSG101 and CD81 content of sEV pellets was determined by Western blotting (B). Values are mean ± SEM, n=7 (A) or n=4 (B) independent experiments, data were analysed by one-way ANOVA. **(C-E)** Parental (P) cells were incubated with the indicated concentrations of buthionine sulphoximine (BSO) for 48 hr. Following this cell conditioned medium was collected for isolation of sEVs by differential centrifugation (E) and the cells lysed for determination of intracellular metabolites by LC-MS (C, D). The heatmap (C) displays levels of the indicated metabolites expressed as log_2_ fold change (normalised to cell number) relative to untreated (P) cells, n=1 (3 technical replicates/condition), and levels of oxidized (GSSG) and reduced (GSH) glutathione are expressed as fold change relative to untreated cells, 3 technical replicates/condition. In the graphs in (D), values are mean ± SD, n=1 (3 technical repeats are shown). The CD63 and TSG101 content of sEV pellets from untreated or BSO treated (1.25μM, 1.88μM, 2.5μM) P cells was determined by Western blotting. CD63 levels in EV pellets were quantified as for figure 5A. Values represent mean ± SEM, n=4 independent experiments (coloured dots), data were analysed by Friedman ANOVA with Dunn’s multiple comparison. **(F, G)** Parental (P), fat pad (FP) or micrometastatic (M) cells were cultured as for figure 2A and levels of intracellular lipidic metabolites determined using LC-MS-based metabolomics. Data are expressed as a volcano plot (F) showing the mean differences (M minus P (M-P); x-axis) between the peak areas of lipids identified in M and P cells. The dotted line represents the p-value (y-axis) above which all the indicated lipid classes display significant differences across the conditions, n=3 independent experiments, data were analysed by paired t-test. The classes of lipids which were detected at significantly different (p<0.033) levels between M and P cells are denoted with coloured dots. Colour-coding for the lipid classes is: cholesterol esters (CE); sphingomyelin/ceramide (SM/Cer); PE, phosphatidylethanolamine (PE); phosphatidylcholine (PC); plasmanyl/plasmenyl phosphatidylcholine (Plas.PC); Bis (monoacylglycerol) phosphate (BMP). Log_2_-fold differences peak areas of the indicated lipid classes (normalised to cell number) between FP and P (FP minus P; blue bars), M and P (M minus P; red bars) are displayed in the graph in (G), n=3 independent experiments, data were analysed using two-way ANOVA with Tukey’s multiple comparison test.

### Micrometastatic cells release sEVs via a neutral sphingomyelinase2-dependent and Rab27-independent mechanism

Altered lipid metabolism is now recognised to be an acquired feature of cancer cells which enables them to meet anabolic and catabolic needs in the face of rapid cell growth ^30^. Moreover, accumulation of lipids, increased uptake of fatty acids and upregulation of genes encoding fatty acid biosynthesis or fatty acid transporters have been shown to enhance migratory and invasive traits of cancer cells and to be associated with metastatic progression in many cancers ^31–33^. To gain a picture of lipid classes which might differ between cells from micrometastases (M) and primary tumours (P & FP), we performed an untargeted lipidomic analysis. This clearly identified two main lipid classes – cholesterol esters (CE) and sphingomyelin/ceramides (SM/Cer) – as being increased in micrometastatic (M) cells with respect to those from primary tumours (P & FP) (Fig. 5F, G). As sphingomyelin metabolism and the production of ceramides have been previously shown to promote the budding of intralumenal vesicles within mutivesicular endosomes, and thereby influence sEV biogenesis and membrane cargo sorting ^34^, we decided to focus on studying the alterations in this class of lipids. Immunofluorescence (using a pan-ceramide antibody) confirmed that ceramide levels were generally elevated in micrometastatic cells and that these lipids appeared to be localised to the plasma membrane and to puncta distributed throughout the cytoplasmic space (Fig. 6A). We then performed a targeted lipidomic analysis - focussing on ceramides and sphingomyelins - which clearly identified four distinct ceramide species whose levels were elevated in micrometastatic cells (M) with respect to their primary tumour counterparts (P & FP) (Fig. 6B). As hydrolysis of sphingomyelins to ceramides contributes to sEV production ^34^, and these lipids are present in sEVs, we also profiled the ceramide content of sEVs. This indicated that two of the species which were elevated in micrometastatic cells (Cer 35:2:2 & Cer 35:3:2) were significantly enriched in sEVs released by these cells (Fig. 6C). We then hypothesised that increased sEV-mediated release of these ceramide species by micrometastatic cells might indicate that these lipids may be used as metastatic biomarkers. To test this, we compared the levels of ceramide species in blood plasma from healthy subjects and from patients with metastatic breast cancer. This indicated that three of the ceramide species (Cer 35:2:2, Cer 35:3:2, & Cer 40:4:2) which were enriched in micrometastatic cells from the MMTV-*PyMT* mouse model (two of which were also elevated in sEV pellets from these cells) were present at significantly increased levels in the plasma of metastatic breast cancer patients (Fig. 6D). This indicates the possibility that release of sEVs rich in certain ceramides from metastatic cells may reflect the presence of metastases in humans with breast cancer.

**Fig. 6.**
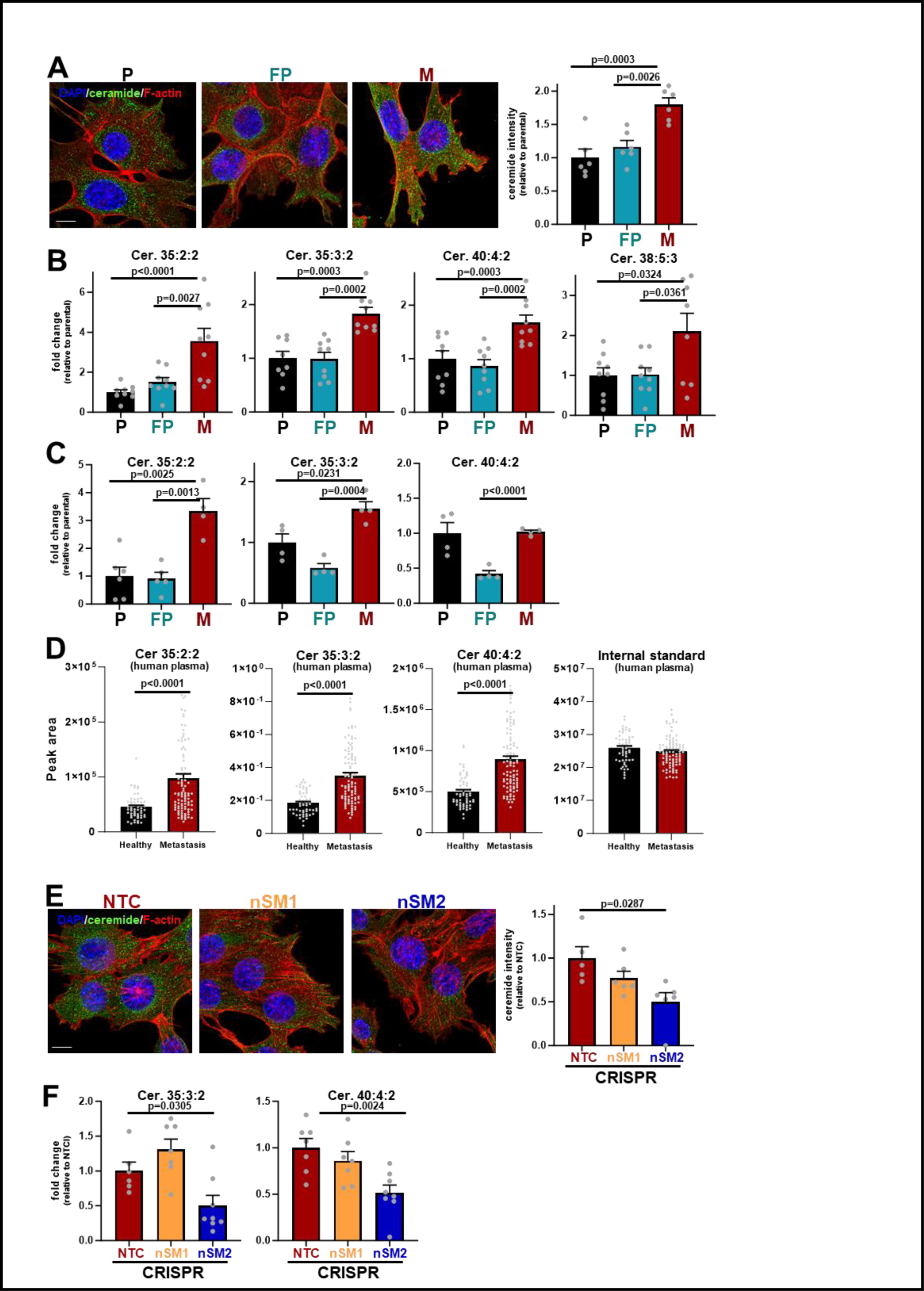
Micrometastatic cells and their sEVs are enriched in ceramide species. **(A)** Parental (P), fat pad (FP) and micrometastatic (M) cells were cultured for 48 hr on glass bottomed dishes and fixed using paraformaldehyde. Ceramides (green), F-actin (phalloidin; red) and nuclei (DAPI; blue) were visualised by immunofluorescence, bar is 10μm. ImageJ was used to quantify the mean intensity of ceramide (sum of z-stacks, 10 stacks/field of view). n>5, data were analysed by ordinary one-way ANOVA. **(B)** The intracellular levels of ceramide species in P, FP and M cells were determined as for figure 5(F, G). Ceramide species found to be present at significantly different levels between M and FP or P cells were normalised to cell number and expressed as fold change relative to (P) cells, values are mean ± SEM, data were analysed using one-way ANOVA with Tukey’s comparison test, n=3 independent experiments (each dot is a technical replicate). **(C)** P, FP, and M cells were cultured for 48 hr. Conditioned media were harvested from these cultures and sEVs purified using differential centrifugation. LC-MS-based lipidomics was used to determine the levels of the indicated ceramide species in the sEV pellets, values are mean ± SEM, n=4 independent experiments, data are analysed by one-way ANOVA. **(D)** LC-MS metabolomics was used to determine levels of the indicated ceramide species in plasma collected from metastatic breast cancer patients (n=96) and matched healthy volunteers (n=55). An internal standard (Splash II, Avanti) was spiked in the lipid extraction buffer, and was analysed to account for technical variation among samples (right graph). Values are mean ±SEM, data were analysed by unpaired Student’s t-test. **(E-F)** M cells were transduced with a lentiviral vectors bearing guide RNAs against non-targeting sequences (NTC) or recognising sequences in nSMase1 (nSM1) or nSMase2 (nSM2). Cells were plated onto glass-bottom dishes and fixed 48h post-seeding. Ceramides were determined using immunofluorescence as for (A) or LC-MS as for (B). Values are mean ± SEM, n=5 in (E) and n=3 independent experiments in (F), data were analysed by one-way ANOVA with Tukey’s comparison test.

Ceramides are formed from sphingomyelins by the hydrolytic removal of a phosphocholine group by neutral sphingomyelinases ^34^. Furthermore, because ceramides promote inward budding of endosomal membranes, neutral sphingomyelinases contribute to multivesicular body (MVB) formation and, in turn, EV release. We used CRISPR to delete the genes for the two major forms of neutral sphingomyelinases (Fig. S4A, B) and found that knockout of neutral sphingomyelinase-2 (nSM2), but not neutral sphingomyelinase-1 (nSM1), significantly reduced ceramide levels in micrometastatic cells (Fig. 6E,F). We then proceeded to determine the neutral sphingomyelinase-dependence of sEV release by cells from primary tumours and micrometastases. This indicated that sEV release from micrometastatic (M) cells was significantly reduced by nSM2 (but not nSM1) knockout, whereas sEV release by cells from primary tumours (P & FP) was independent of nSM1 and nSM2 (Fig. 7A, B; S5A, B). Moreover, treatment of primary tumour cells (P) with 1.88 µM BSO (to judiciously decrease glutathione levels) drove sEV release that was nSM2-dependent (Fig. 7C).

**Fig. 7.**
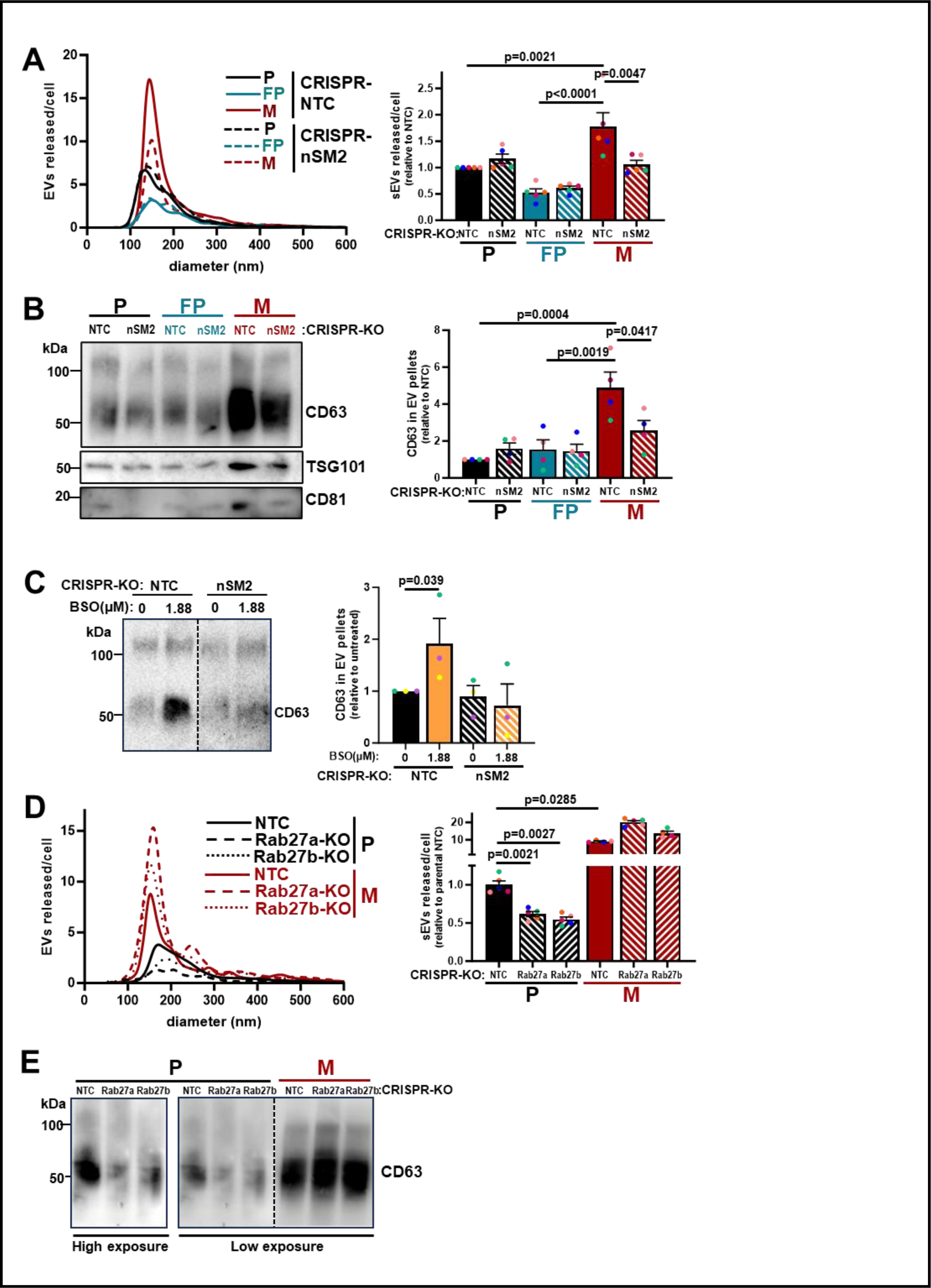
Micrometastatic cells release sEVs via a nSMase2-dependent and Rab27-independent mechanism. Parental (P), fat-pad (FP) or micrometastatic (M) cells were transduced with a lentiviral vector bearing guide RNAs recognising non-targeting sequences (NTC) or sequences in nSMase2 (nSM2), Rab27a or Rab27b. [See figure S4 for confirmation of nSMase and *Rab27* gene deletion]. sEVs were purified from conditioned medium collected over a 48 hr period analysed using nanoparticle tracking (A, D) as for figure 5A, and levels of CD63 in sEV pellets were determined by Western blotting (B, C, E). In (C) P cells transduced with non-targeting (NTC) lentiviral vectors or those targeting nSMase-2 (nSM2) were incubated with the buthionine sulphoximine (BSO; 1.88 µM) during the 48 hr EV collection period. Values are mean ± SEM, data were analysed by one-way ANOVA, n=5 (A, D), n=4 (B) n=3 (C) independent experiments

The Rab27 GTPases are also key regulators of EV release ^35^. We, therefore, used CRISPR to delete the genes for Rab27a and Rab27b in both micrometastatic and primary tumour cells (Fig. S4C) and measured sEV release into their conditioned media. As anticipated, deletion of Rab27a and Rab27b strongly opposed release of sEVs from cells from primary tumours (P) (Fig. 7D, E). However, sEV release from micrometastatic cells (M) was not opposed by deletion of Rab27s and was even increased following CRISPR-mediated knockout of Rab27a (Fig. 7D, E). Taken together, these data indicate that, when mammary cancer cells begin to colonise the lung, they may upregulate ceramide levels to enable a switch from Rab27-dependent/nSM2-independent sEV production to a mechanism of sEV release which is nSM2-dependent, but independent from Rab27s.

### The switch to nSM2-dependent sEV release enables micrometastatic cells to generate pro-invasive microenvironments

Upregulated ceramide levels and the consequent switch to nSM2-dependent sEV release prompted us to investigate the role of this enzyme in invasiveness of micrometastatic cells. Deletion of nSM2 did not influence transmigration toward a gradient of serum and fibronectin (not shown), indicating that cell-autonomous migratory behaviour depended on neither nSM2 nor EV release from micrometastatic cells. However, despite their intrinsically migratory capabilities, nSM2 knockout micrometastatic cells displayed reduced ability to move through ‘organotypic’ collagen plugs (Fig. 8A). As the microenvironment of organotypic plugs is strongly influenced by fibroblasts, this indicated the possibility that nSM2-dependent sEV release enables communication between micrometastatic cells and fibroblasts to permit cancer cell invasiveness. To test this, we pre-treated fibroblasts with sEVs from control or nSM2 knockout cells prior to introducing them into collagen plugs ^16^. sEV-treated fibroblasts were then allowed to precondition collagen plugs for 4-5 days. Control or nSM2 knockout micrometastatic cells were then plated onto, and allowed to invade into, preconditioned plugs for a further 7 days (Fig. 8B). This indicated that when collagen plugs were pre-conditioned with sEVs from control (but not nSM2-knockout) micrometastatic cells, this restored the ability of nSM2-knockout cells to invade into the organotypic microenvironment (Fig. 8C).

**Fig. 8.**
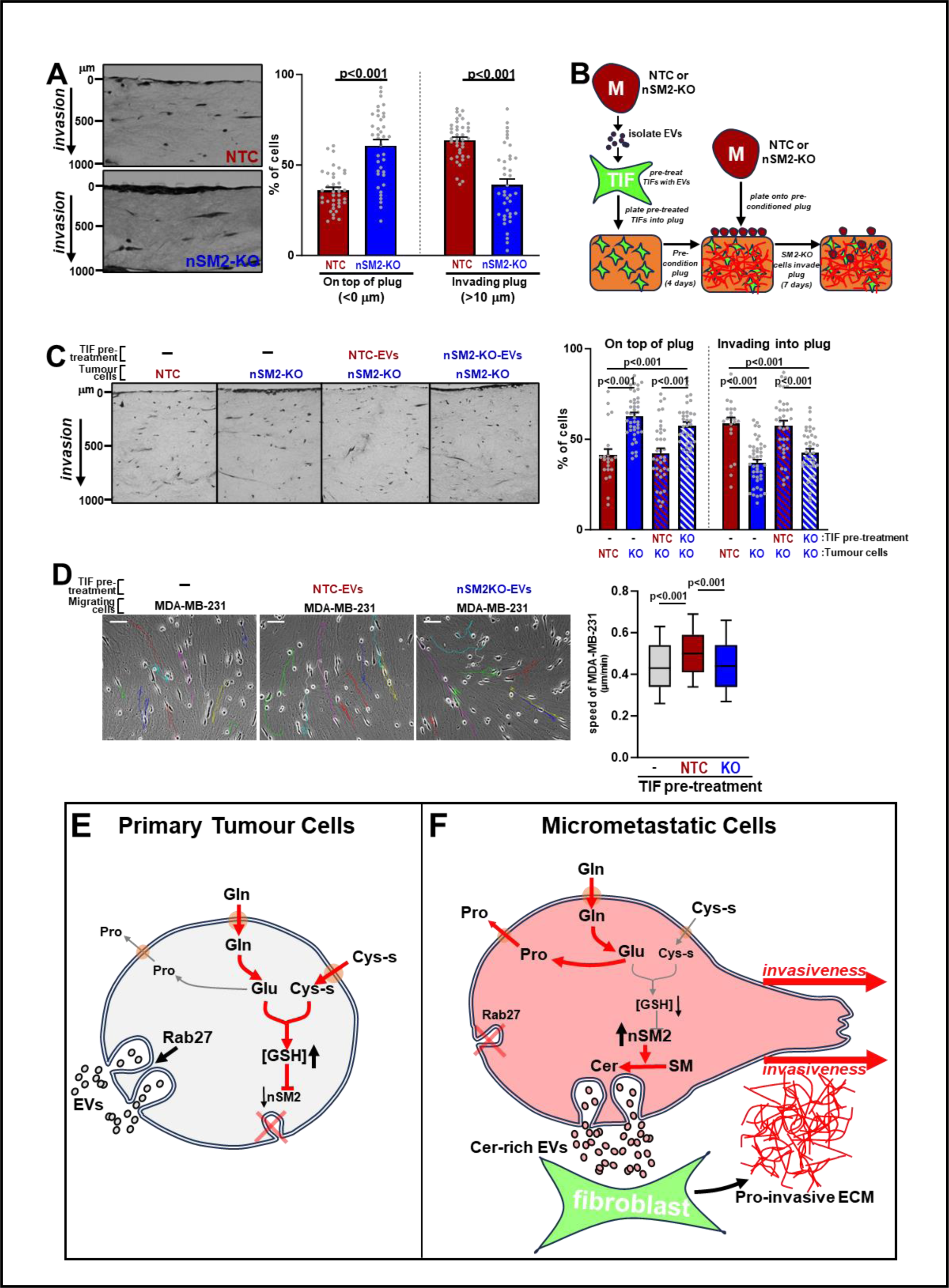
nSMase2-dependent sEV release favours generation of a pro-invasive microenvironment. **(A)** Micrometastatic (M) cells transduced with lentiviruses bearing guide RNAs recognising non-targeting sequences (NTC) or sequences in nSMase2 (nSM2) were plated onto fibroblast pre-conditioned plugs and invasion into these determined as for figure 1E. Values are mean ± SEM, 2 plugs/condition, n=39-47 fields of view, data were analysed by mixed effects ANOVA with Tukey’s multiple comparison test. **(B)** Schematic depiction of protocol for determining sEVs’ influence on the invasive microenvironment of organotypic collagen plugs. sEVs released by control (NTC) or nSMase-2 knockout (nSM2-KO) M cells were incubated with telomerase immortalised fibroblasts (TIFs) for 72 hr. sEV-treated TIFs were then allowed to pre-condition organotypic plugs of rat tail collagen for 4 days. NTC or nSM2-KO M cells were then plated onto pre-conditioned plugs and allowed to invade for 7 days. **(C)** Control (NTC) or nSMase-2 knockout (nSM2-KO) M cells were plated onto organotypic collagen plugs that had been pre-conditioned as outlined in (B). Invasion was quantified as for figure 1E. Values are mean ± SEM, n>22 fields of view/condition (n=2 independent experiments) data were analysed using mixed effects ANOVA with Tukey’s multiple comparison test. **(D)** TIFs were incubated with sEVs from control (NTC) or nSMase-2 knockout (nSM2-KO) M cells for 72 hr or were left untreated. sEV pre-treated TIFs were then allowed to generate extracellular matrix (ECM) for 7 days, which was then de-cellularised. MDA-MB-231 breast cancer cells were plated onto de-cellularised ECMs and timelapse microscopy (over 16h) followed by cell tracking (ImageJ) were employed to measure their migration (n>100 cells, 2 independent experiments). Coloured lines indicate representative tracks of individual migrating cells (left panels). Bar, 100 μm. Box and whisker plots are 10-90 percentile, line display median. Data were analysed by Kruskal-Wallis test with Dunn’s multiple comparisons test. **(E-F)** Schematic summary of how metabolic control of sEV biogenesis in cells from primary mammary tumours and lung micrometastases may influence ECM microenvironments. Cells from primary tumours synthesise glutathione using glutamine-derived carbons and release sEVs in a Rab27-dependent/nSMase2-independent manner (E). In micrometastatic cells, more glutamine-derived carbons are used for proline production leading to decreased glutathione synthesis. This, in combination with increased nSMase2-dependent ceramide production, promotes release of sEVs in a Rab27-independent/nSMase2-dependent way, which encourages fibroblasts to generate a more invasive microenvironment (F) (Cys-s, cystine).

This result led us to propose that sEVs released in a nSM2-dependent manner from micrometastatic cells may modify the microenvironment of the organotypic plugs by influencing how the fibroblasts deposit and/or remodel the ECM ^16^. To test this, we pre-treated the TIFs with sEVs from control or nSM2 knockout micrometastatic cells (as for Fig. 8B) but, instead of introducing them into collagen plugs, we allowed them to deposit ECM in 2D culture for 7 days. We then de-cellularised the ECM and measured migration speed of MDA-MB-231 mammary cancer cells plated onto this. Pre-treatment of TIFs with sEVs from control, but not nSM2 knockout, micrometastatic cells increased the migration speed of MDA-MB-231 cells subsequently plated onto ECM deposited by these fibroblasts (Fig. 8D). Taken together, these data indicate that when disseminated mammary cancer cells move to the lung and form micrometastases, they alter their metabolism in a way that favours nSM2-dependent/Rab27-independent production of sEVs. sEVs released via this nSM2-dependent route then influence ECM deposition by fibroblasts to foster a microenvironment which may contribute to subsequent metastatic colonisation of the lung.

## Discussion

Metabolic reprogramming enables cancer cells to overcome challenges encountered on the path to metastasis and may enhance their potential to seed distant organs. We show here that cells from mammary cancer lung micrometastases have rewired their metabolism in a way that generates an invasive microenvironment. The metabolic adaptations of lung micrometastases are manifest in the re-routing of glutamine-derived carbons toward proline production (and secretion) at the expense of glutathione synthesis – likely owing to reduced xCT and GCL levels, and increased nSM2-dependent synthesis of ceramides. The combination of decreased glutathione synthesis and increased ceramide levels promote sEV release via a Rab27-independent/nSM2-dependent route which, in turn, influences ECM deposition/remodelling by fibroblasts to foster an invasive microenvironment.

Evidence is accumulating that alterations to proline metabolism can drive breast cancer metastasis ^12, 26, 36^, and we concur that proline metabolism is indeed re-wired during early metastasis. Micrometastatic cells from the lungs of mice bearing MMTV-*PyMT* tumours increase proline production and release it into the medium at the expense of glutathione synthesis. Moreover, because pharmacological inhibition of proline synthesis ^27^ can, at least in part, restore glutathione synthesis to micrometastatic cells this indicates that the need to increase proline production may be a metabolic priority during early metastatic seeding. Our data indicate that most of the proline synthesised by micrometastatic cells is not retained in the cells but exported into the medium, suggesting that a role for these cells may be to increase proline levels in the extracellular microenvironment during early metastasis. As ECM components, particularly collagens, have very high proline content it is, therefore, possible that proline exported from micrometastatic cells may be used by other cells in the lung microenvironment to synthesise collagen-rich ECM. Indeed, a recent study has described how myofibroblastic-type carcinoma-associated fibroblasts (myCAFs) re-wire their metabolism to favour synthesis of proline via PYCR which is then used for collagen production to drive tumour progression and aggression ^26^. Such a requirement for metabolic support to ECM production during early metastatic seeding has been highlighted by a study describing how breast cancer cells can generate metastatic niches in the lung by increasing production of αKG to activate the hydroxylation of prolines in collagens ^12^. In addition to supporting ECM production, evidence is accumulating that regulation of proline synthesis plays a key role in maintaining redox balance in metabolically stressed cells. Proline synthesis via PYCR can lead to NAD(P)H consumption, thus providing a route to regeneration of NAD(P)+ without the cell needing to activate oxidative phosphorylation and the consequent generation of damaging reactive oxygen species (ROS) ^37, 38^. Indeed, metabolic flux analysis indicates that the micrometastatic cells of the present study do not display significantly elevated oxygen consumption (not shown), despite increased flux of glutamine-derived carbons into the Krebs cycle. This raises the possibility that micrometastatic cells exploit proline synthesis in a manner that allows sufficient Krebbs cycle activity to maintain anaplerosis, but without passing high energy electrons to the electron transport chain. Thus, micrometastatic cells may avoid excessive ROS generation which would constitute a risk in the highly oxidative environment of the lung.

Increased proline synthesis may reduce ROS generation, but the fact that this occurs at the expense of glutathione production still presents micrometastatic cells with a redox challenge. Although there have been long-running controversies over the role of antioxidants in cancer therapy, it is now established that glutathione synthesis and maintenance of a reducing cellular environment is necessary for both primary tumour initiation and growth ^39^. Consistently, several studies now demonstrate that inhibition of glutathione synthesis – typically by targeting xCT – increases cancer cell death and inhibits tumour growth ^40, 41^. The reasons for this are multifarious and likely include roles for the glutathione/thioredoxin system in inhibiting ferroptosis (and other pathways driving cell death) and the ability of antioxidants to prolong oncogene-induced senescence during early tumorigenesis by reducing oxidative stress. Furthermore, metastatic outgrowth – a later event in metastasis - also relies heavily on glutathione synthesis, as highlighted by a recent study describing how cell lines from established colon cancer (macro-)metastases are particularly sensitive to inhibition of cysteine import via xCT ^42^. Nevertheless, although a level of glutathione synthesis is clearly a requirement for growth of primary tumours and of established metastases, this does not preclude transient adoption of metabolic states characterised by low glutathione levels during early metastatic seeding. Indeed, a recent study has demonstrated that, although glutathione production is required for tumour initiation in mammary carcinogenesis, it becomes dispensable later in disease progression ^39^. These authors showed that inhibition of glutathione synthesis decreases tumour burden only when administered prior to primary tumour onset. Inhibition of glutathione synthesis following tumour onset evokes alternative antioxidant responses enabling glutathione-depleted cells to thrive despite oxidative insults. Another recent study shows that high expression levels of xCT promote primary tumour growth but suppress metastasis ^43^. The mechanistic interpretation offered by these authors is that metastasising cancer cells with high xCT activity are particularly susceptible to oxidative stress because this increases intracellular cystine. Subsequent conversion of intracellular cystine to cysteine – a redox reaction necessary for glutathione synthesis - mops up intracellular NADPH rendering the metastasising cells sensitive to disulphide stress and death. Our finding that micrometastatic cells cultured from the lungs of mice bearing MMTV-*PyMT* tumours display reduced xCT and GCL also suggests that moderate expression levels of xCT – and, therefore, glutathione synthesis - may be necessary during the early stages of metastatic seeding because of the need to upregulate proline synthesis. Increased proline synthesis and its release not only helps to maintain redox balance, but also offers the possibility of contributing to ECM synthesis by other cells in lung microenvironment to condition the early metastatic niche.

Decreased glutathione, in combination with increased ceramide levels, contributes to switching off Rab27-dependent sEV release and encourages micrometastatic cells to release sEVs via a nSM2-dependent pathway. nSM2-dependent production of sEVs supports deposition of pro-invasive ECM by fibroblasts, and it is interesting to compare this with other situations through which altered landscapes in tumour cells can, via sEV release, foster invasiveness in other cells. Metabolic stress in tumour cells promotes PINK1-dependent packaging of mtDNA into EVs that are released via the Rab27 pathway, and mtDNA-containing EVs alter invasive behaviour of other tumour cells by activating TLR9 signalling ^19^. However, nSM2-dependent (unlike Rab27-dependent) EV production is not associated with release of mtDNA (not shown). Thus, although this and the present study both describe mechanisms through which altered tumour cell metabolism can communicate invasiveness (via sEVs) to other cells, the mechanisms mediating this must be distinct. Other studies from our laboratory have shown that acquisition of gain-of-function mutations in p53 alters the podocalyxin content of sEVs ^16^. This, in turn, alters integrin trafficking in fibroblasts to encourage them to deposit a more pro-invasive ECM. However, we have measured α5β1 integrin recycling and found this not to differ between fibroblasts treated with sEVs from control and nSM2-CRISPR micrometastatic cells. Thus, further work will be necessary to determine how sEVs released via a nSM2-dependent pathway influence ECM deposition by fibroblasts, and it will be interesting to evaluate the role of lipidic EV cargoes, such as ceramides and cholesterol esters, in this process.

Taken together, our study has elucidated the metabolic reprogramming which occurs in mammary cancer cells as they seed lung metastases, how this influences the cellular machinery controlling sEV production and how these EVs, in turn, can foster a pro-invasive microenvironment. Thus, we provide mechanistic insights that further our understanding of how changes in cellular metabolic programs can coordinate membrane trafficking processes to influence the lung microenvironment in a way that is likely to render this organ more fertile for subsequent metastatic outgrowth. We believe that the mechanistic understanding outlined in the present study will enable further exploration of the metabolic vulnerabilities that may be targeted to oppose metastatic outgrowth of breast cancer in the lung.

## Materials and Methods

### MMTV-*PyMT* model of mammary cancer

All mice carrying a mouse mammary tumour virus (MMTV) promoter-driven polyoma middle T (*PyMT*) transgene were backcrossed >20 generations in FVB/N background. The MMTV-*PyMT* mice have been described previously ^44^. MMTV-*PyMT* mice (Jackson Laboratory, ME, USA) were housed in individual-ventilated cages with environmental enrichment, in a barrier facility (12-hour light/dark cycle). Monitoring of mice for tumour development was performed two to three times per week. Tumour growth was monitored by calliper measurement with clinical endpoint any tumour reaching 15 mm diameter when mice were humanely euthanised by Schedule 1 methods. Metastatic burden (number of metastases, metastatic index) in the lungs was assessed by H&E staining. Metastatic burden was scored by slicing the formalin-fixed paraffin embedded lungs into sections and staining them with 3 H&E section numbers 5, 10 and 15 (each 5 sections apart) were then scored for presence of metastases. All work was carried out with ethical approval from CRUK Scotland Institute and University of Glasgow under the revised Animal (Scientific Procedures) Act 1986 and the EU Directive 2010/63/EU (PP6345023). All animal experiments were performed in accordance with relevant guidelines and regulations (3Rs).

### Mammary fat pad transplantation and resection

For orthotopic transplantation experiments, 8-to-10-week-old female FVB/N mice were obtained from Charles River (UK). 0.5 x10^6^ cells generated from MMTV-*PyMT* tumours (either P or P’; see generation of *PyMT* cell lines below) in 50μl PBS/Matrigel (1:1) were transplanted into the fourth mammary fat pad (inguinal) of FVB mice ^21^. Following recovery from surgery, mice were monitored for tumour development at least three times per week. Once tumour was palpable, growth was assessed by calliper measurement, and tumour was surgically removed when the diameter reached 8-10mm. The tumour was kept in ice-cold PBS prior to downstream processing for cell line generation (see below). After surgical resection, mice were closely monitored (three or more times per week) for clinical signs of metastasis, which included weight loss, altered respiration or abdominal distension. Animals were humanely euthanised by Schedule 1 methods, if they displayed two or more moderate signs (e.g. weight loss and moderate piloerection). At clinical endpoint, lungs were harvested, kept in ice-cold PBS prior to further processing for generation of cell lines. Analgesia was used prior and after all surgical procedures which were carried out under anaesthesia.

### Generation of cell lines

Mammary tumours (FP and FP’ lines) or lungs (M and M’ lines) were minced using sterile surgical scalpels into a fine paste, and the finely chopped tissue was added to culture media (DMEM), followed by centrifugation at 1000rpm for 5 minutes at room temperature (RT). The cell pellet was resuspended in DMEM, filtered using a 70μm cell strainer (to remove remaining clumps) and added to a 10-cm dish. The media was supplemented with 10% FBS, 2mM glutamine, 100U/ml penicillin streptomycin,

20 ng/ml hEGF, 10 μg/ml insulin and 0.25μg/ml fungizone and cultured at 37°C/ 5% CO_2_ in a humidified incubator. Prior to mincing, resected lungs were inspected for using a dissection microscope. Any micrometastases that were visible were dissected from the lung tissue at this stage and minced and cultured independently. The macrometastatic cells (maM’) used in this study originated as a macrometastasis dissected from a lung belonging to a mouse implanted with the P’ parental MMTV-*PyMT* line. To establish an immortalised mammary tumour cell line, cells were passaged (once reached around 90% confluence) for at least five generations and maintained in culture for a maximum of twenty passages. The same protocol was used for the generation of metastatic cell lines, with the exception that following centrifugation, the resuspended cell pellet was seeded in 6 well plates. For cell culture growth and maintenance, cells were kept in the aforementioned conditions (without fungizone supplementation after the fifth passage) and passaged every two days. All cells were regularly tested for mycoplasma contamination.

### Immunofluorescence

Cells were seeded (3×10^4^ cells/dish) in 35 mm glass bottomed dishes and incubated at 37°C/ 5% CO2 for 40 hours and fixed with 4% PFA (in PBS) at room temperature for 15 minutes. Cells were permeabilised with 0.1% Triton-X100 (in PBS) at room temperature for 15 minutes, and blocked in 1% BSA (in PBS) for 1 hour. Cells were then incubated with an antibody recognising ceramides (Enzo, ALX-804-196; dilution 1:100) in blocking solution at 4°C overnight, followed by secondary antibody (Alexa Fluor 488 anti-Mouse IgG, ThermoFisher (A-21202), dilution 1:1000) and phalloidin stain (1:1000 dilution in PBS). Cells were then mounted using Vectashield (containing DAPI; Vectashield Laboratories) and visualised by confocal microscopy using an Airyscan super-resolution microscope. ImageJ software was used to quantify the mean intensity of ceramide stain (sum of z-stacks, 10 stacks/field of view).

### In situ hybridisation

Formalin fixed paraffin embedded (FFPE) tissues were obtained from MMTV-*PyMT* primary mammary tumours and matched metastatic lungs. FFPE tissue was cut into 4µm sections which were then heated to 60°C for 2 hours. H&E staining was performed on a Leica autostainer, where sections were dewaxed, taken through graded alcohols and stained with Haem Z (CellPath) for 13 minutes. Sections were then washed in tap water, differentiated in 1% acid alcohol, washed and the nuclei blued in Scotts tap water substitute (in-house). After washing, sections were placed in Putt’s Eosin (in-house) for 3 minutes. In situ hybridisation (ISH) detection for Slc7a11 (xCT) mRNA was performed using RNAScope 2.5 LSx (Brown) detection kit (Bio-Techne) on a Leica Bond Rx autostainer strictly according to the manufacturer’s instructions. To complete H&E and ISH staining, sections were rinsed in tap water, dehydrated through graded ethanols and placed in xylene. The stained sections were mounted in xylene using DPX mountant (CellPath). Following staining, slides were scanned at 20× magnification using NanoZoomer NDP scanner (Hamamatsu), and HALO software (Indica Labs) was used to quantify the average optical density of the *Slc7a11* stain in sections of primary mammary tumours and corresponding lung metastases.

### Western blotting

Proteins were resolved in 4-12% or 10% pre-cast NuPAGE Bis-Tris gels run using MOPs running buffer at 120V. Resolved proteins were transferred to PVDF membrane in NuPAGE transfer buffer at 120V for 90 minutes. Following blocking (in 4% milk) membranes were incubated with primary antibodies diluted in 1% milk, overnight at 4°C while gentle agitation was applied. Primary antibodies were: antiCD63, Abcam (ab217345) dilution 1:1000; anti-CD81, Santa-Cruz (sc-166029) dilution 1:1000; anti-TSG101, GeneTex (GTX70255), dilution 1:1000; anti-vinculin, Sigma (MAB3574-C), dilution 1:5000. Following incubation with appropriate HRP-conjugated secondary antibodies, protein bands were visualised using enhanced chemiluminescence (SuperSignal West Femto) for 1 minute in a Bio-Rad ChemiDoc imaging system. Densitometry analysis of CD63, quantification was performed at non-saturating exposures, while applying the same background settings across the samples, using Image Lab software (Bio-Rad).

### qPCR

Reverse transcription quantitative real-time PCR (RT-qPCR) reactions were performed in 96-well plates (BioRad CFX platform) by using SyBr green in a total reaction volume of 10μl per well. Each reaction contained a 1/20 dilution of cDNA template, 1X PerfeCTa SYBR Green Fast mix, 0.5μM forward and reverse primers and samples were plated in duplicate.

Primer sequences were as follows:

ARPP P0 (forward) GCACTGGAAGTCCAACTACTTC

ARPP P0 (reverse) TGAGGTCCTCCTTGGTGAACAC

*Gclc* (forward) CCACAAGAGCTCATCCTCCCT

*Gclc* (reverse) GGGTCGGATGGTTGGGGTT

*PyMT* (forward) CTGCTACTGCACCCAGACAA

*PyMT* (reverse) GCAGGTAAGAGGCATTCTGC

SLC7A11 (xCT) (forward) CTTTGTTGCCCTCTCCTGCTTC

SLC7A11 (xCT) (reverse) CAGAGGAGTGTGCTTGTGGACA

Smpd2 (neutral sphingomyelinase-1) (forward) CATCCCCTACCTGAGCAAAC

Smpd2 (neutral sphingomyelinase-1) (reverse) CCAGGAGAGCCAGATCAAAGTT

Smpd3 (neutral sphingomyelinase-2) (forward) TGGTGGTGTTTGACGTCATCT

Smpd3 (neutral sphingomyelinase-2) (reverse) GCGAGTAAAGAGCGAGTGCT

Standard curves were generated using serially diluted cDNA from pooled samples to assess reaction efficiency, and no template controls (without cDNA) were also included for each target gene to check for potential primer dimer formation. Reactions were run on the BioRad C1000 thermal cycler using a 3 step protocol, during which cDNA was denatured for 3 minutes at 95°C, followed by 40 cycles of denaturation at 95°C for 20 seconds, annealing at 60°C for 20 seconds, and extension at 72°C for 20 seconds, with a final extension step of 72°C for 5 minutes and a melting curve from 65°C to 95°C in 0.5°C increments. Data acquisition and analysis were performed in CFX Manager Software, and the ΔΔCt method was used to compare gene expression levels between different cell lines. In all analyses the ARPP P0 mRNA was used as internal control.

### Cell proliferation

Cells were seeded at 2×10^4^ and 3×10^4^ cells per well in 6-well culture plates and incubated at 37°C/ 5% CO_2_ in a humidified incubator for 6 hours to adhere. Plates were transferred to a tissue culture incubator equipped with IncuCyte ZOOM live-cell imaging system, and images were acquired (10× objective) every 2 hours over a period of 120 hours (5 days). At the end of the incubation, images were analysed using the Incucyte Base Analysis Software, and confluence was measured using the same phase segmentation parameters across all *PyMT*-derived cell lines. Confluence was calculated as a % of the phase image area covered by cells (3 technical replicates or fields of view/condition) corresponding to individual time points, and these values (sixty time points) were plotted over the culture period to generate the growth curves for each cell line. The time elapsed (t_1/2_) until half of the phase image area per field of view was covered by cells (50% confluence) was also plotted in these graphs.

### Generation of nSMase and Rab27 CRISPR cells

Guide RNA (gRNA) sequences targeting neutral sphingomyelinases 1 & 2 and Rab27s a and b were cloned into a lentiCRISPR vector (Addgene (52961) ^45, 46^. gRNAs were as follows:

Rab27a (forward) CACCG CCAGGAGCATCTCAATCGCG;

Rab27a (reverse) AAACCGCGATTGAGATG CTCCTGGC;

Rab27b (forward) CACCGGCTGCGCTTGTTTCGAAGTA;

Rab27b (reverse) AAACTACTTCGAAACAAGCGCAGCC;

neutral sphingomyelinase-1 (forward) CACCGCGCCCTATGTTTGCTCAGGT;

neutral sphingomyelinase-1 (reverse) AAACACCTGAGCAAACATAGGGCGC;

neutral sphingomyelinase-2 (forward) CACCGGTGGTGTTTGACGTCATCTG;

neutral sphingomyelinase-2 (reverse) AAACCAGATGACGTCAAACACCACC.

Then, to produce lentivirus encoding Cas9, along with the desired gRNAs, HEK 293T cells were used as the host packaging cell line. Virus-containing HEK 293T supernatant was then filtered through a 0.45μm PTFE filter membrane, polybrene supplemented to a final concentration of 4μg/ml and the medium incubated with recipient P, FP, or M cells overnight at 37°C/ 5% CO_2_. Virus transfected cells were then selected using puromycin (2μg/ml), the antibiotic selection medium being replenished every 2 days, and the cells were passaged according to their confluence to generate stable cell lines. 7 passages post infection, cells were harvested for qPCR analysis of nSMases and Western blot for Rab27s.

### Metabolite extraction, calculation of metabolite exchange and metabolic tracing

Cells were seeded in 6-well plates (3 technical replicates/cell line) in full culture medium (2ml) at 1.5×10^4^ cells per well. After 24 hours (day 1), 2ml of culture medium was added to each well to prevent nutrient exhaustion. On day 2, culture medium was changed to 2ml or 7ml for medium metabolite extractions (calculation of exchange rates) or intracellular metabolite extractions respectively. For the metabolic tracing experiments, DMEM in which ^13^C_5_-labelled glutamine was substituted for unlabelled glutamine was deployed at this stage. The pyrroline-5-carboxylate reductase inhibitor (PYCRi), prepared as described in ^27^ was added at this stage at the indicated concentrations. The volumes of culture medium used enabled measurement of exchange rates of nutrients with slow flux, while detecting intracellularly metabolites that are rapidly consumed. In parallel, at this time point (day 2) cells were harvested for determination of total cell number per condition. After 24 hours (day 3), media and intracellular metabolites were extracted. For extraction of metabolites from culture media, 20μl of medium was added to 980 μl of an ice-cold polar extraction solution (methanol, acetonitrile, and water in a 5:3:2 ratio), and the samples were then shaken in a thermomixer at 1400rpm for 10 minutes (4°C). Metabolites were extracted from serum and plasma using the same approach as for cell culture medium. For cell extracts, cells were quickly washed three times with ice-cold PBS, followed by extraction with 600 μl of ice-cold extraction solution for 5 min at 4°C, whilst gentle agitation was applied. To remove insoluble material, media and cell extracts were centrifuged at 16000g for 10 minutes, and the supernatant was transferred to glass vials, which were stored at -80 °C prior to LC-MS analysis.

Lipids were extracted from cells using 600μl, or from sEV pellets using 50 µl, of a solution composed of butanol and methanol (1:1), supplemented with Splash II standard. Lipid cell extracts were centrifuged at 16000g for 10 minutes to remove insoluble material, and the supernatant was transferred to glass vials, which were stored at -80 °C prior to LC-MS analysis.

To allow estimation of consumption/secretion rates of nutrients and metabolites, data normalisation was achieved by counting the total number (#) of cells on days 2 and 3, and the exchange rate per day for each metabolite was calculated according to the following [inline1], where Δmetabolite= (detected peak area spent medium-detected peak area cell-free medium), and whether the x value is positive or negative indicates that the metabolite is secreted or consumed by the cells respectively ^47^.

### LC-MS metabolomics and data analysis

All polar metabolites were analysed by Liquid chromatography-Mass Spectrometry (LC-MS) as previously described in ^48^.

#### Polar metabolites

Metabolites from the biological extracts (5 μl) were separated using a ZIC-pHILIC guard and analytical column (SeQuant; 150 mm by 2.1 mm, 5 μm; Merck) on an Ultimate 3000 high-performance liquid chromatography (HPLC) system (Thermo Fisher Scientific). Chromatographic separation was performed using a 15 min linear gradient from 20% ammonium carbonate [20 mM (pH 9.2)] and 80% acetonitrile (ACN), to 20% ACN at a constant flow rate of 200 μl/min. The column temperature was constant at 45°C. A Q Exactive orbitrap mass spectrometer (Thermo Fisher Scientific) equipped with electrospray ionization was coupled to the HPLC system, polarity switching mode was used with a resolution (RES) of 70,000 at 200 mass/charge ratio (m/z), to enable both positive and negative ions to be detected across a mass range of 75 to 1000 m/z (automatic gain control target 1e6, and maximal injection time 250 ms). Polar metabolomics analysis was performed using Tracefinder (version 4.1, Thermo Fisher Scientific). Extracted ion chromatograms were generated for each compound and its isotopologues using the m/z of the singly charged ions (XIC, +/-5 ppm) and the RT (+/-2 min) from our in-house metabolite library, which was generated using reference standards on the same LC-MS method. Thiols were preserved as previously described in ^24^. Standards were derivatised and analysed with the same methodology to generate optimised XIC parameters. All peak areas were normalised to total cell number e.g. peak area per million cells.

#### Lipdomics (targeted analysis)

Lipid analysis was performed using an Ultimate 3000 HPLC (Thermo Fisher Scientific) coupled to a Q-Exactive Orbitrap mass spectrometer (Thermo Fisher Scientific). Lipids were separated on an Acquity UPLC CSH C18 column (100 × 2.1 mm; 1.7 µm; Waters Corporation) maintained at 60°C. The mobile phases consisted of 60:40 ACN: H_2_O with 10 mM ammonium formate, 0.1% formic acid and 5 µM of phosphoric acid (A) and 90:10 IPA:ACN with 10 mM ammonium formate, 0.1% formic acid and 5 µM phosphoric acid (B). The gradient was as follows: 0–2 min 30% (B); 2–8 min 50% (B); 8–15 min 99% (B); 15–16 min 99% (B); and 16–17 min 30% (B). Sample temperature was maintained at 6°C in the autosampler and 5 µL of sample were injected into the LC-MS instrument. Thermo Q-Exactive Orbitrap MS instrument was operated in both positive and negative polarities, over the following mass range 240–1,200 m/z (positive) and 240–1,600 (negative) at RES 70,000. Data-dependent fragmentation (dd-MS/MS) was carried out for each polarity. Feature detection and peak alignment from .Raw files were performed using Compound Discoverer 3.1 (Thermo Fisher Scientific, Waltham). Files were also converted to .mgf format using MSConvert software (ProteoWizard) and searched against the LipiDex_ULCFA database using LipiDex software ^49^. Data pre-processing, filtering and basic statistics were performed using Perseus software ^50^ version 1.6.2.2, after importing the final results table from LipiDex.

#### Lipdomics (untargeted analysis of ceramides)

A lipidomic platform was established to identify and measure the ceramides present in the biological extracts using tandem mass spectrometry. The analysis was performed using an Ultimate 3000 HPLC (Thermo Fisher Scientific) coupled to a TSQ Altis Triple quadrupole mass spectrometer. The method used the same chromatographic conditions applied in the untargeted lipidomic analysis. MS was performed employing SRM lists generated using LipidCreator version 1.2.1 ^51^. SRM lists included all possible fragments for species with saturated or unsaturated fatty acid of chain length (C16–C20) based on preliminary experiments on abundance of ceramide species of variable chain lengths. After peak deconvolution and integration of the chromatograms using TF software (version 4.1, Thermo Fisher Scientific). Initially, 1579 peaks were detected, species were considered only if they showed fragments for both the sphingoid units and the expected acyl chains, all at the same retention time. This strategy was applied for structural identification of ceramides. Data cleaning and filtering were all done using R software. Species were excluded if detected in blanks or were of ≥ 40% missingness, with relative standard error (RSE) of > 20%. This approach narrowed down the number of detected species across the groups to 12 species. Fold chain analysis and 1.5 was selected as an arbitrary threshold in the metastatic compared to parental & FPs which left only four species found to be significantly different in the metastatic group versus control.

### Organotypic collagen plug invasion assay

Organotypic plugs were generated from rat tail-derived collagen I, as previously described ^22^. Briefly, 1×10^6^ confluent TIFs were mixed with collagen I (2mg/ml) under neutral or slightly acidic conditions (MEM, approximate pH=7.2 by using 0.22M NaOH) and the collagen/fibroblast matrix was allowed to contract for approximately 4 days, until it fitted in a 24-well culture dish. 3×10^3^ Parental (P), fat pad (FP) and micrometastatic (M) cells were seeded on top of these plugs (in duplicates) and cultured for 2 days. Plugs were then transferred on top of a metal grid and cultured in full medium by creating an air-liquid interface (invasion day 0) for 7 days. At the end of the incubation time, plugs were fixed in 4% paraformaldehyde before paraffin embedding (proper orientation of cross sections), and 4μm sections were then cut and stained (H&E). Single cell tracking was performed manually using ImageJ software under enhanced brightness/contrast settings (black and white mode) and a distance value, along with an angle value (representing the orientation relative to the invasion baseline; negative angle: cell entering the plug, positive angle: cell residing on top of the plug) were assigned to each cell per field of view. Images were acquired using a brightfield microscope and raw data were exported in batches (.ome.tiff) using the Zeiss Zen lite software (blue edition).

### Production of TIF-derived ECM

TIFs (2×10^5^ cells/well) that had previously been exposed to sEVs for 72 hr were seeded in 6-well plates coated with gelatin and cells were incubated at 37°C/5% CO_2_. Once cells reached confluence, the medium was changed to full medium supplemented with 50μg/ml ascorbic acid, which were refreshed every other day for seven days. Extracellular matrices (ECMs) were denuded by incubation with a Triton-containing buffer (2 minutes at RT), residual DNA was digested with DNase I (10μg/ml in PBS containing calcium and magnesium; D-PBS), and denuded ECMs stored at 4°C. Before use, CDMs were washed twice in D-PBS and incubated with full medium for 30 minutes at 37°C/5% CO_2_. MDA-MB-231 cells were seeded at 0.8×10^5^ cells per well and imaged in time-lapse for 16 hr using an inverted phase-contrast microscope (Axiovert S100, Carl Zeiss, 10× objective), while kept at 37°C/ 5% CO_2_ (6 fields of view per well were imaged every 10 minutes for 16 hours). After completion of image acquisition, movies were analysed using ImageJ software to determine cell migration speed.

### Transwell migration assay

These assays were performed in Boyden Chambers (Corning, Merck, 3422; Transwell Costar, 6.5 mm diameter, 8 μm pore size) according to the manufacturer’s instructions ^52^. Briefly, to assess migration the lower faces of the inserts were coated with 2 μg*/*ml fibronectin (Merck, F0895). Subsequently, cells (1 x 10^5^ cells/well) were resuspended in serum-free medium and added to the upper chamber, while 10% FBS-containing medium was added to the lower chamber. 2 hr later, inserts were removed, and the upper face of each insert was washed to remove cells that did not transmigrate. Cells that had transmigrated to the lower chamber were stained with 0.1% crystal violet in 2% ethanol and visualised using an inverted microscope.

### sEV collection and nanoparticle tracking analysis

EV-free medium was prepared by overnight (16 hours) ultracentrifugation of 5% FBS-containing DMEM at 100,000g (4°C) and further filtration of the collected supernatant using a 0.2µm PES filter unit. 0.3× 10^6^ cells were then seeded in 15cm dishes (three 15cm dishes/condition), and cultured in full medium for 16 hours at 37°C/5% CO_2_. Cells were then washed twice with PBS, medium was changed to EV-free medium (15ml/dish) and cultured for 48 hours at 37°C/5% CO_2_. This conditioned medium was then collected and subjected to sequential centrifugation steps to remove live cells (300g for 10 minutes), dead cells (2,000g for 10 minutes) and finally to remove cell debris and larger lipid membrane particles, including the majority of microvesicles (10,000g for 20 minutes). sEVs were then pelleted in thin wall polypropylene tubes after ultracentrifugation at 100,000g for 70 minutes using a SW32 rotor (Beckman coulter). The sEV pellet was then washed in filtered PBS (0.2µm PES) and subjected to a final ultracentrifugation step at 100,000g for 70 minutes. All centrifugation steps were performed at 4°C. At the end of the last spin, the supernatant was carefully aspirated by tilting the tube, EVs were re-suspended in 150μl PBS, which was previously filtered using a Whatman Anotop filter (0.02µm) and stored at 4°C. Nanoparticle tracking analysis was performed using a NanoSight LM10 instrument (Malvern Panalytical) with a high sensitivity camera according to the manufacturer’s instructions. Isolated EVs were diluted 1:50-1:200 in PBS (filtered through 0.02µm). Diluted EV samples were then flushed through the chamber until vesicles were visible on the camera by using a 1ml syringe. The focus and gain settings were optimised for each run, and kept consistent across the samples, which were injected into the flow cell and 5 recordings of 60 seconds each were acquired. The chamber was washed with ethanol and deionised water between samples to ensure no residual particles remained. The data were analysed using the NTA3.1 analysis software and average of technical replicates were plotted per experiment following normalisation to the total number of EV-releasing cells and the dilution applied per condition.

### Statistical analyses

Statistical analyses were performed with GraphPad Prism 9 on all relevant experiments using unpaired t-tests to compare two groups with normal data distribution. ANOVA tests (one-way, two-way or repeated measures) were used to compare more than two groups if the data were normally distributed, while a Kruskal-Wallis test was performed for not normally distributed datasets. A Friedman test (nonparametric repeated measures ANOVA) was used when no assumptions were made for the data distribution. Statistical significance is annotated in the figures (p– values are shown on the figures, with p<0.05 considered significant) and the associated tests are indicated in each figure legend.

## Acknowledgments

We would like to thank core staff in the Biological Services Unit, the Beatson Advanced Imaging Resource (BAIR), Molecular Services, Histology Facility and Central Services (CRUK-Scotland Institute) for their support which facilitated the work described in this manuscript. This paper was critically reviewed by Catherine Winchester (CRUK Scotland Institute).

## Author contributions

Experimental design: MG; AVC; ES; LM; IM; ST; DS; KB; JCN; CJC

Investigation: MG; AVC; ES; LM; ED; NR; SD; LP; IM; ST; DS; KB; JCN; CJC

Reagents: EK; SZ; JM; CJ

Funding acquisition: JCN; CJC; IM; KB

Writing – original draft: JCN; MG; CJC

Writing – review & editing: MG; AVC; ES; LM; IM; ST; DS; KB; JCN; CJC; SZ; DS; ES

## Data and materials availability

The data supporting the findings of this study are available within the article and its supplementary information files and from the corresponding author upon request.

## Supplemental material

**Fig. S1.**
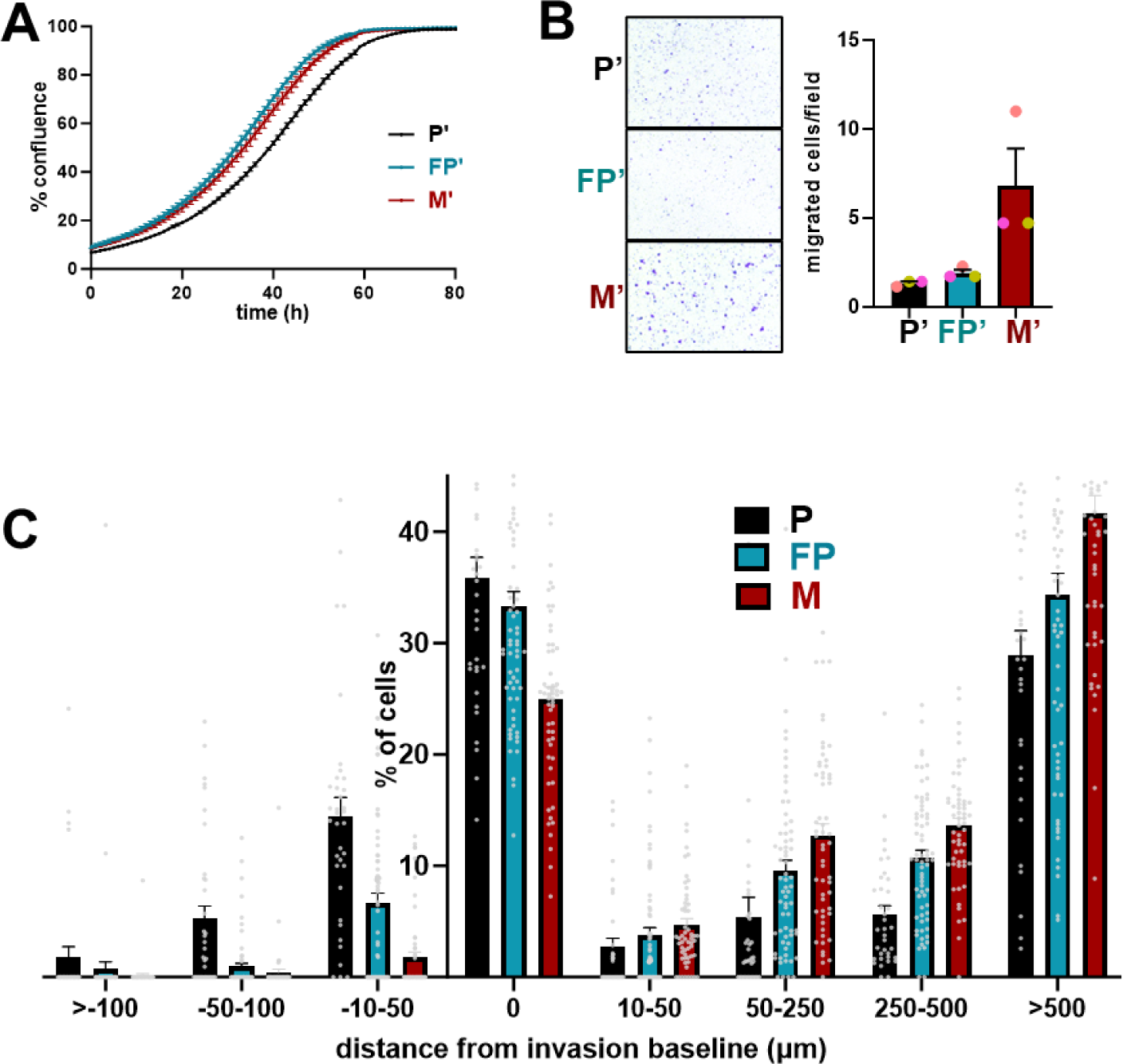
Growth and invasiveness of cells from lung micrometastases and their primary tumour counterparts. The protocol described in figure 1A was used to obtain a duplicate set of MMTV-*PyMT* derived cells lines. These are termed parental’ (P’), fat pad’ (FP’) and micrometastatic’ (M’) cells. **(A, B)** The growth (A) and transmigration (B) of P’, FP’ or M’ cells was determined as for figure 1B & C. **(C)** The distribution of (P), (FP) and (M) cells (expressed as %) at a range of depths of invasion into collagen organotypic plugs was measured by Image J. A negative value in distance from the invasion baseline indicates that cells remain on top of the collagen plug and do not invade. These data relate to those displayed in Fig. 1E.

**Fig. S2.**
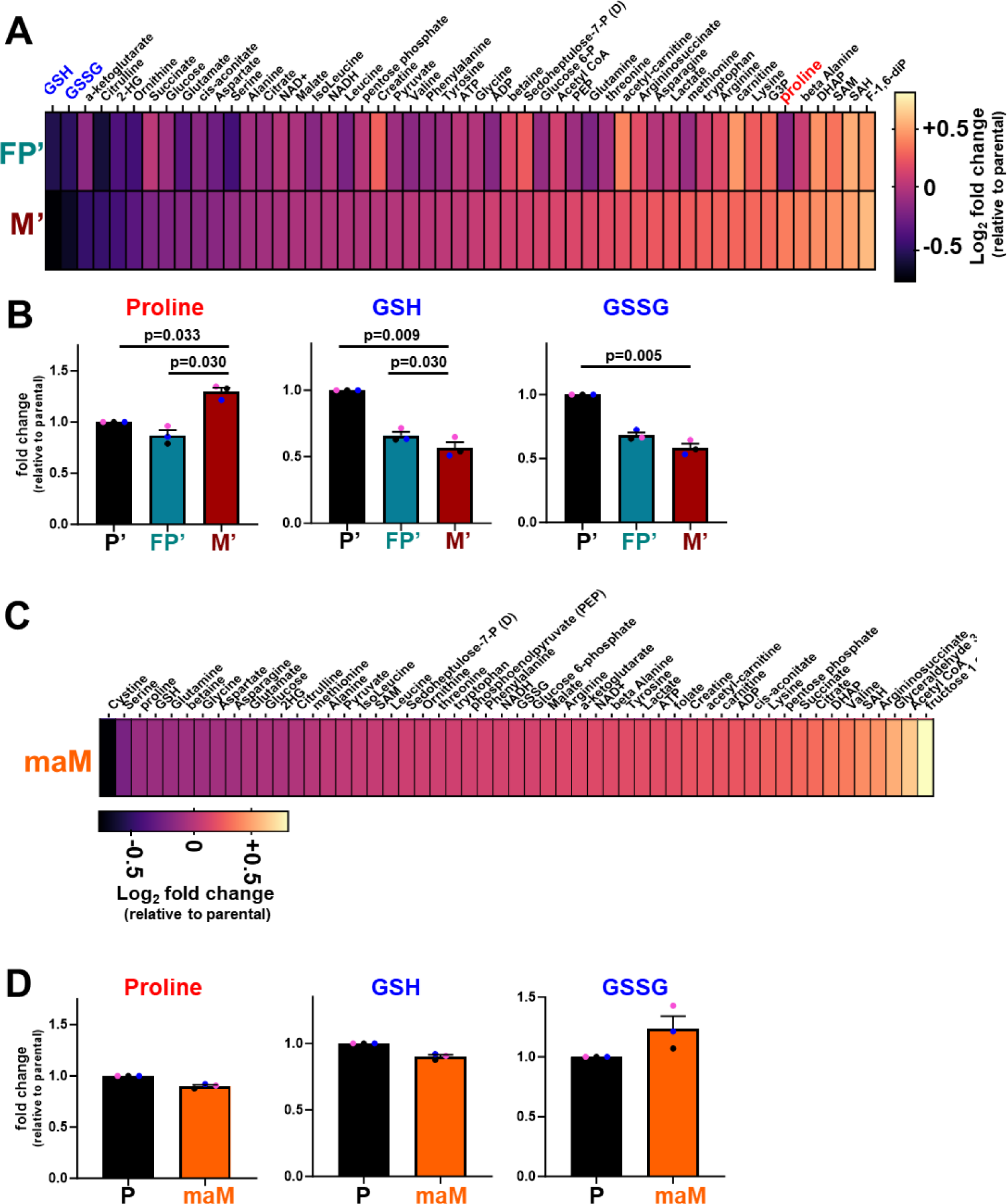
Comparison of metabolite levels of cells from micrometastases and frank metastases with their primary tumour counterparts. The abundance of intracellular metabolites in fat-pad’ (FP’) micrometastatic’ (M’) (A, B) and frank macrometastatic (maM’) (C, D) cells was determined using LC-MS and expressed relative to levels of the same metabolites in P’ cells. For the heatmaps (A, B) values are log2-fold changes, and for the graphs (B, D) values are mean fold-change ± SEM, statistics are repeated measures one-way ANOVA with Tukey’s multiple comparison test, n=5 independent biological replicates.

**Fig. S3.**
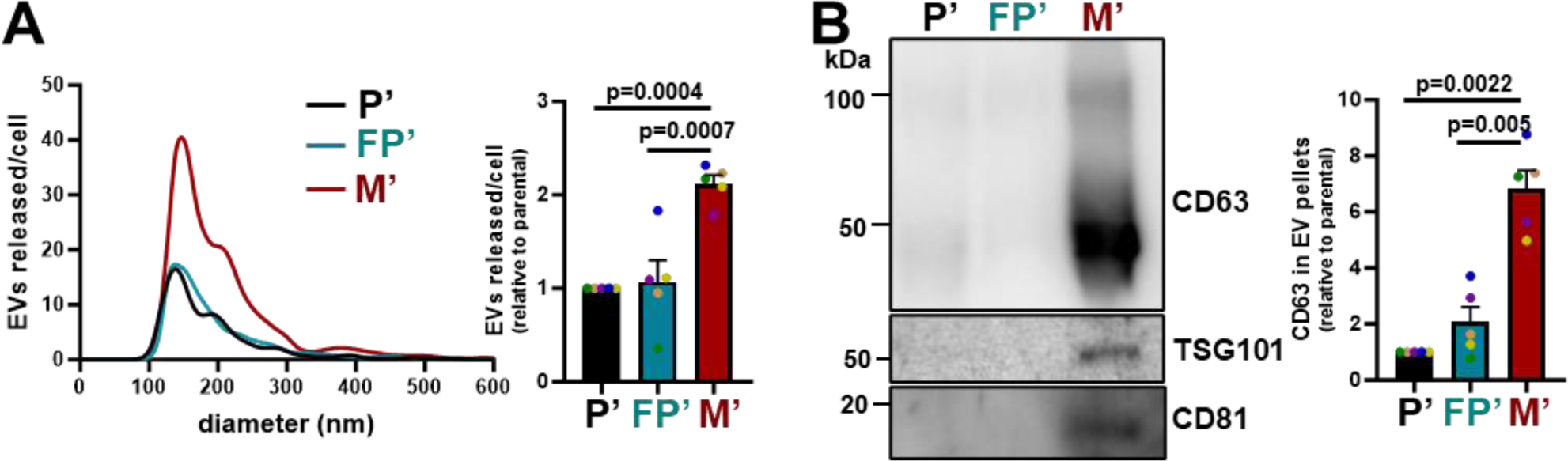
sEV release from micrometastatic cells. sEVs were purified from the conditioned media of parental (P’), fat pad (FP’) and micrometastatic (M’) cells, as in fig. 5A, B. Size distribution and number of EVs were analysed by nanoparticle tracking (A), and CD63, TSG101 and CD81 content of sEV pellets was determined by western blot (B). Total particle counts and sample loading were normalized to the number of EV-releasing cells. Values are mean ± SEM, n=5 independent experiments, one-way ANOVA, p-values are shown on the graph.

**Fig. S4.**
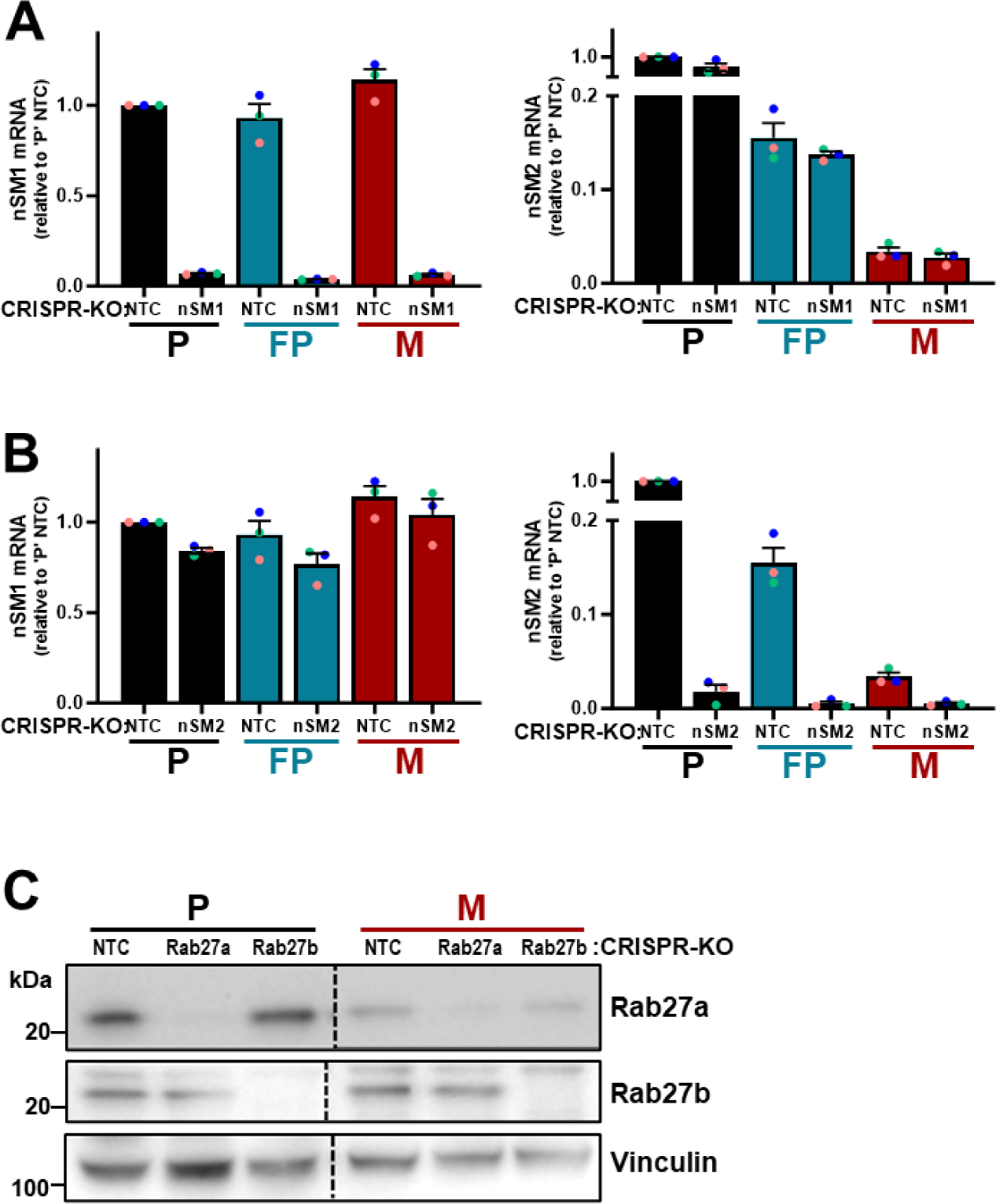
Validation of CRISPR deletion of nSMases and Rab27s. **(A, B)** Parental (P), fat pad (FP) and micrometastatic (M) cells in which nSMase1 (nSM1) or nSMase2 (nSM2) had been disrupted by CRISPR were lysed. Levels of mRNA encoding nSM1 (A) or nSM2 (B) were determined using qPCR. All data were normalised to ARPP P0 and presented relative to expression in parental NTC cells, values are mean ± SEM, n=3 independent repeats. **(C)** Western blotting was used to determine levels of Rab27a and Rab27b in parental (P) and micrometastatic (M) cells in which the genes for the Rab GTPases had been disrupted using CRISPR. Vinculin is used as a loading control.

**Fig. S5.**
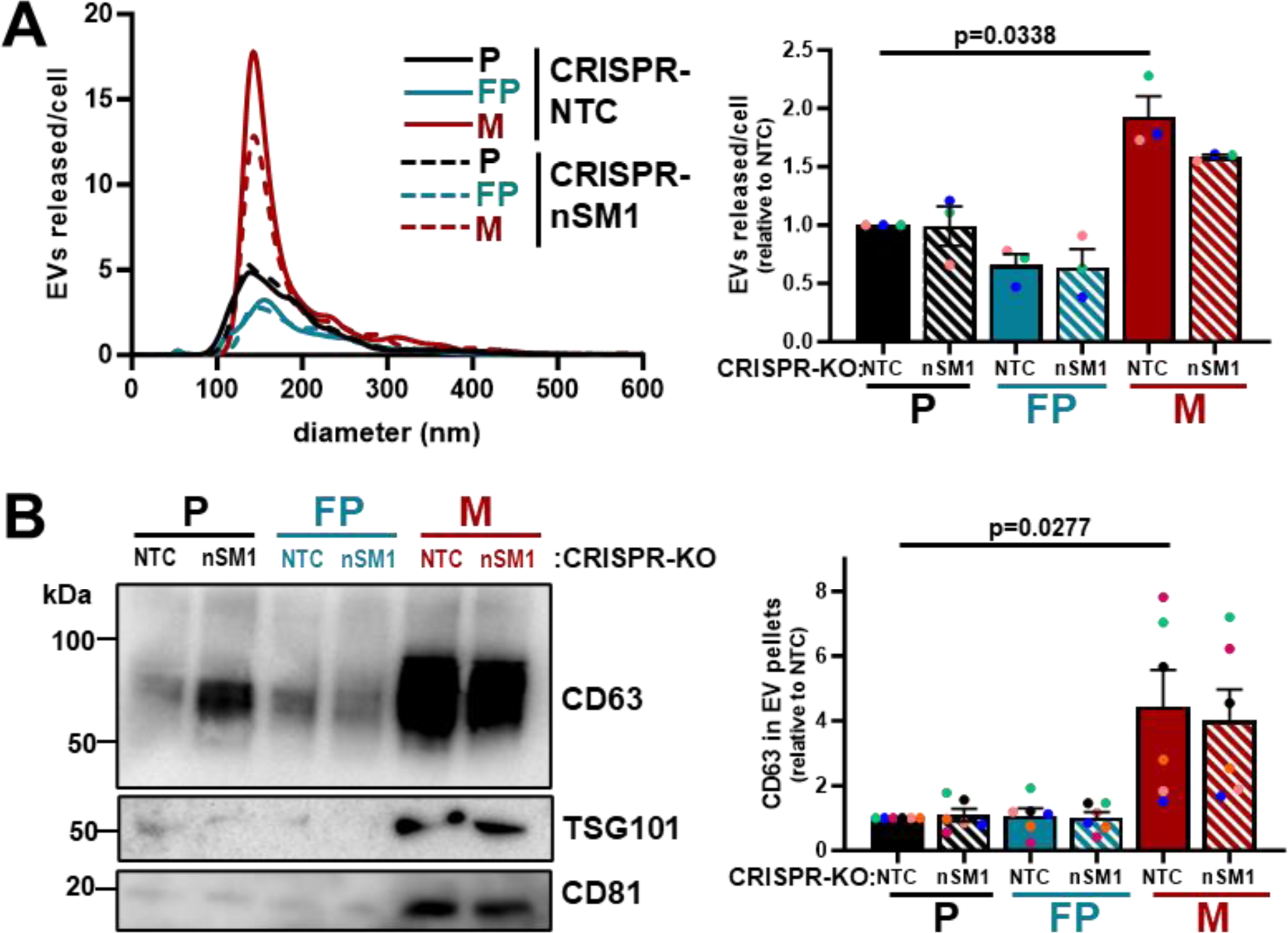
sEV release from nSMase-1 knockout cells. Parental (P), fat-pad (FP) or micrometastatic (M) cells were transduced with a lentiviral vector bearing guide RNAs recognising non-targeting sequences (CRISPR-NTC) or sequences targeting nSMase1 (CRISPR-nSM1). [See figure S4 for confirmation of nSM1 deletion]. EVs were purified from conditioned medium collected over a 48 hr period and analysed using nanoparticle tracking (A) as for figure 5A, and levels of CD63 in sEV pellets were determined by Western blotting as for figure 5B. Values are mean ± SEM, one-way ANOVA, n=3 (A), n=6 (B) independent experiments

